# A competition-protection balance explains the evolution of resistance within simple microbial communities

**DOI:** 10.64898/2026.05.20.726537

**Authors:** Massimo Amicone, Adriana Espinosa-Cantú, Gabriela Petrungaro, Tobias Bollenbach, Sara Mitri

## Abstract

Stressful environments can pose a threat to microbial populations, but resistant individuals can emerge and avoid extinction. Adaptation to stress is classically studied in isolated microbial species, ignoring ecological interactions, a key component of natural ecosystems. A growing body of experimental work has shown that community context can affect resistance evolution due to a large variety of mechanisms. Here we set out to identify the minimal components needed to predict the likelihood of acquiring resistance in a focal species embedded within a simple community. To achieve this, we developed a mathematical model based on evolutionary rescue theory and validated it with two experimental systems: *Escherichia coli* evolving on exposure to the antibiotic nitrofurantoin alone or with one of 14 bacterial isolates from urinary tract infections, and *Microbacterium liquefaciens* evolving in ampicillin alone or with ampicillin-degrading *Comamonas testosteroni*. One key factor that emerged from our analyses – the relative strength of competition versus protection – could explain whether a focal species is more or less likely to evolve resistance in the presence of a partner species. While competition always hinders the emergence of resistance, protection can rescue the focal species in two ways: (i) ecological rescue, when the partner species completely removes the antibiotic and favors the survival of the susceptible population, or (ii) evolutionary rescue, when the partner only lowers antibiotic concentrations and favors the emergence of resistant variants, a previously overlooked evolutionary consequence of detoxification. Overall, by integrating theory and experiments, we propose a framework that clarifies how ecological interactions favor or hinder the evolution of resistance to antibiotics or potentially other stressors.

**Significance:** Bacteria can rapidly adapt to resist stressors, such as antibiotics. While resistance evolution in single populations or species is well understood, it remains unclear how ecological interactions with other species influence this process. We develop a mathematical framework to predict what interactions should favor resistance evolution and validate it with two sets of experiments where bacteria adapt to antibiotics in small communities. Our work demonstrates that interactions with other species shape the probability of evolving resistance in a predictable way, determined by the balance between competition and protection against the stressor. By identifying the key factors that drive these dynamics, our work helps explain how bacteria adapt to environmental challenges within species-rich ecosystems.

## Introduction

Microorganisms influence the health of every ecosystem on Earth, from the human body to the open ocean. In each of these ecosystems, tens to thousands of different microbial species can assemble into relatively stable communities. Environmental challenges, such as global warming or antimicrobial treatments, can disrupt these stable ecosystems, leading to downstream effects that are difficult to predict [Cavicchioli et al., 2019]. Within microbial communities, interactions among species are pervasive and large populations can lead to rapid adaptation [Mitri and Foster, 2013, Neher, 2013]. Ecological and evolutionary dynamics play out over similar timescales [Garud, 2021, Good and Rosenfeld, 2023], and our understanding of how microbial communities respond to environmental disturbances remains limited.

One way populations respond to environmental challenges is through evolutionary rescue: the emergence of a resistant genotype which can spread and take over the declining population, otherwise destined to extinction [Bell, 2017]. In bacteria, a classical example of evolutionary rescue is antimicrobial resistance (AMR). When faced with antibiotics, bacterial populations can survive by evolving resistant phenotypes, but interactions with other species can influence this adaptive process. A growing list of experiments (both *in vitro* and *in vivo*) demonstrate that the ecological context can shape bacterial responses to antibiotics [Quinn et al., 2022, Munch et al., 2023], and that this can happen through a variety of different mechanisms [DeWit et al., 2022]. Yet, a deeper understanding of AMR evolution within communities requires not only biological evidence, but also theoretical frameworks that can simplify what appears to be a complex network of many possible ecological and evolutionary mechanisms and integrate them into a unifying quantitative framework [DeVos et al., 2018, Bottery et al., 2021].

Here we start with a simple question: when do other microbial species favor or hinder the emergence of resistance in a focal species? To reduce the complexity of the many potentially contributing factors, we describe the emergence of resistance through evolutionary rescue processes and develop a theoretical framework to compare the likelihood of surviving a stressful environment in isolation or in the presence of another species. We find that a key factor can explain when ecological interactions should favor resistance evolution: the balance between stress-independent decrease of population size (competition) and the increase of tolerance to stress (protection).

We then validated model predictions using two sets of evolution experiments involving species pairs.

First, we followed the dynamics of an *Escherichia coli* strain in mono- and co-culture with different urinary tract infection (UTI) bacterial isolates [Croxall et al., 2011, DeVos et al., 2017]. UTIs are primarily caused by pathogenic *E. coli*, but in certain patient groups, such as immunocompromised, post-menopausal, or elderly individuals, they can involve multiple bacterial strains and species with a high incidence of resistance to drugs [Croxall et al., 2011, Kline and Lewis, 2016]. This makes the UTI isolates an ideal experimental model to validate our framework of resistance within communities, while better understanding a clinically relevant system. Here, we demonstrate how interactions shape the evolution of resistance to nitrofurantoin, and predict the observed rescue probabilities through the competitive and protective effects identified with the model.

To further validate our framework, the second set of experiments involved two environmental bacteria (*Microbacterium liquefaciens Ml*, and *Comamonas testosteroni Ct*), whose ecological interactions were previously described in our lab [Piccardi et al., 2019, DosSantos et al., 2022]. We were able to control the strength of the competitive and protective effects of *Ct* by tuning the medium composition, and to use the model to explain how their interplay shaped the evolutionary response of *Ml* to the antibiotic ampi-cillin.

By combining mathematical models with experimental evolution of bacteria isolated from clinical or environmental settings, we show how the emergence of resistance in simple communities can be explained by the relative contribution, or balance, of competitive and protective interactions.

## Results

### Competition and protection under a two-species evolutionary rescue model

To set up a model that encompasses the diversity of possible mechanisms by which ecological interactions may influence resistance evolution [DeWit et al., 2022], we use the evolutionary rescue framework. Classical evolutionary rescue theory describes the dynamics of a population that upon an environmental change (e.g. exposure to antibiotics) goes extinct, unless a genetically resistant variant emerges and rescues the population, generating the classical U-shaped curves (Fig. **1**a) [Alexander et al., 2014].

**Fig. 1:**
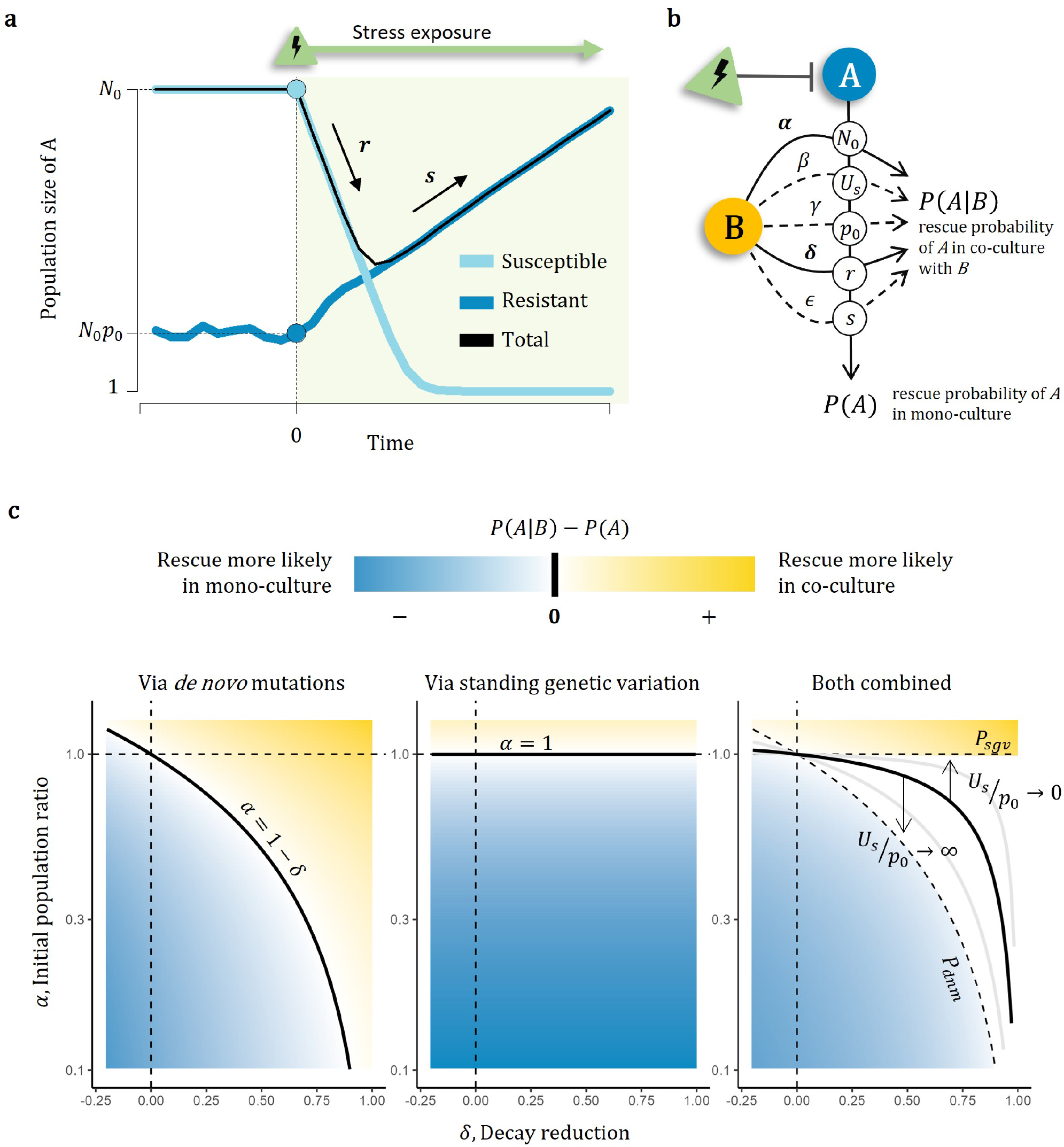
Evolutionary rescue theory and the Competition-Protection balance. **a)** Example of evolutionary rescue dynamics where a susceptible genotype goes extinct upon exposure to a stress (e.g. antibiotic), but the emergence of a resistant type rescues the entire population. **b)** Schematics of the key variables. *N*_0_, *U*_*s*_, *p*_0_, *r* and *s* represent respectively the initial population size, the input of resistant types via mutation, the initial fraction of resistant type, the rate at which the susceptible type decays and the rate at which the resistant type grows. These are the key properties of species *A* determining its rescue probability in mono-culture, *P* (*A*). In co-culture with a second species *B*, the rescue probability of *A* (*P* (*A*| *B*)) will depend on the parameters *α, β, γ, δ, ϵ* representing the effects that a second species *B* has on the previously defined variables. **c)** Assum-ing *β, γ, ϵ* = 1 we show the conditions under which *B* does not affect (black line), promotes (blue region) or hinders (gold region) the rescue probability of *A*, via *de novo* mutations (**c-left**), via standing genetic variation (**c-center)**), or via a combination of the previous (**c-right)**). In the white region, the transition from negative to positive depends on the relative contribution of new mutations (*U*_*s*_) and standing variation (*p*_0_). In this example the solid line for the rightmost plot was obtained with *U*_*s*_*/p*_0_ = 0.001.

We model the dynamics of a focal species *A* under stress: the susceptible genotype (*a*) decays geometrically by *N*_*t*_(*a*) = *N*_*t−*1_(*a*)(1 *− r*), while the resistant genotype (*a*′) grows by *N*_*t*_(*a*′) = *N*_*t−*1_(*a*′)(1 + *s*), where *N*_*t*_(*a*) is the abundance of type *a* at a given time *t*. Taking into account the stochastic nature of replication and mutation, the rescue probability depends on five key parameters: the initial population size (*N*_0_), the initial fraction of the resistant type (*p*_0_), the rate at which the susceptible type decays (*r*), the rate at which the resistant type grows (*s*), and the input of resistant types via mutation (*U*_*s*_). Following Orr and Unckless [Orr and Unckless, 2008, Orr and Unckless, 2014], it is possible to derive an approximation for the probability of rescue via *de novo* mutations (*P*_dnm_), via standing genetic variation (*P*_sgv_), or via both processes combined (*P*_tot_) (*Methods*).

We then ask how these rescue probabilities change under the influence of a second species. Assuming that this second species *B* is resistant to the stress allows us to describe its interactions as constant effects on the global rescue parameters (Fig. **1**b). We consider a scenario where *B* only affects the initial population size of *A* by 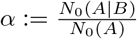, and the decay rate by 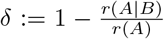. We focused on these because they capture common ecological effects and are the easiest to manipulate experimentally, but the model can be extended to study evolutionary rescue under the influence of all five parameters (*Methods* and *Supplementary materials*).

The regime where *α <* 1 captures interactions whereby *B* reduces the population size of *A*, prior to the environmental change (e.g. due to competition); the regime where 0 *< δ <* 1 captures interactions whereby *B* partially protects *A* (e.g. via detoxification); the extreme case when *δ* = 1 is what we call the *ecological* rescue scenario, whereby the protective effect completely prevents the decay and genetic changes are not needed to rescue the population. Here, the probability of evolutionary rescue is not defined (*Methods*), but mutations have virtually infinite time to emerge and fix.

To predict when *B* should favor or hinder the emergence of resistance in *A*, we derive the competition-protection balance – the conditions under which the probability of rescue in mono-culture equals the probability of rescue in co-culture (*P* (*A*) = *P* (*A* |*B*)) (Fig. **1**c). The balance via *de novo* mutations (*P*_dnm_(*A*) = *P*_dnm_(*A* |*B*)) is achieved when *α* = 1 −*δ* (Fig. **1**c, left). Intuitively, interactions that lower the population size decrease the effective input of mutations (*NU*_*s*_), but interactions that lower the decay rate can counteract this effect by keeping the population alive longer. Thus, the stronger the competition between the two species is, the stronger the protective effect has to be to balance the probability of rescue via *de novo* mutations. Instead, rescue via standing genetic variation does not depend on the decay rate (as we assume that the resistant mutant growth is independent from the susceptible one), such that *P*_sgv_(*A*) = *P*_sgv_(*A*| *B*) simply when *α* = 1 (Fig. **1**c, center).

When both mechanisms are combined (*P*_tot_ =*P*_sgv_ + (1 *− P*_sgv_)*P*_dnm_), the ecological effects *α* and *δ* only provide the boundary conditions: when *α >* 1 we expect *B* to favor rescue of *A*, while when *α <* 1− *δ* we expect *B* to hinder it. For all intermediate cases (1 −*δ < α <* 1) we need to know the source of resistant mutations (*U*_*s*_ and *p*_0_), and the exact conditions that define the competition-protection balance depend on *U*_*s*_*/p*_0_ (Fig. **1**c, right). This result can be explained in terms of resistance cost: Assuming that a resistance mutation is selected against in the absence of the stress, the stronger the fitness cost (*c*), the lower its frequency should be prior to antibiotic exposure. Over long time-scales, the frequency of a costly mutation should stabilize according to the mutation-selection balance 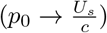 [Johnson, 1999]. In our framework, this means that costly resistance should contribute little to standing genetic variation, and we can approximate the competition-protection balance with the *de novo* mutations case, while non-costly mutations should contribute more to standing variation (Fig. **1**c, right). Because in experimental setups populations are initially isogenic and undergo only a few growth cycles before exposure to stress, when analyzing our experimental dynamics we will assume that the contribution of *de novo* mutations is stronger than that of standing variation.

Taken together, our simple phenomenological model identifies two distinct regimes: one where the partner species favors the evolution of resistance by increasing stress tolerance and/or limiting competition (yellow region of Fig. **1**c), and one where the partner species hinders it (blue region of Fig. **1**c). We next explore the two different regimes and test our model’s predictions through experimental evolution of bacteria exposed to antibiotics.

### Competition with urinary tract infection isolates predicts the emergence of resistance to nitrofurantoin

To test whether the competition-protection balance described above can explain resistance evolution in co-cultures with isolates from UTIs, we set up an *in vitro* experiment where a focal strain (*Escherichia coli* MG1655) undergoes serial passaging in Artificial Urinary Medium (AUM), with or without nitrofurantoin (an antibiotic commonly used to treat UTIs) and with or without a second partner strain (Fig. **2**a). Drug treatment consisted of 8.56 μg/ml nitrofurantoin, a concentration above the minimum inhibitory concentration (MIC) of *E. coli* that can sustain partial growth (Fig. S1). Out of a collection of over 70 natural bacterial isolates [Croxall et al., 2011, DeVos et al., 2017], we selected 14 UTI partner strains that were resistant to the chosen nitrofurantoin concentration, showed diverse interactions with our focal strain, and span the commonly observed phylogeny of polymicrobial UTI iso-lates (*Methods*). We followed the abundance of the focal strain via luminescence over 9 transfers in 15 culture combinations (1 mono-culture and 14 co-cultures), with or without nitrofurantoin (in 4 or 50 independent replicates, respectively) (*Methods*).

**Fig. 2:**
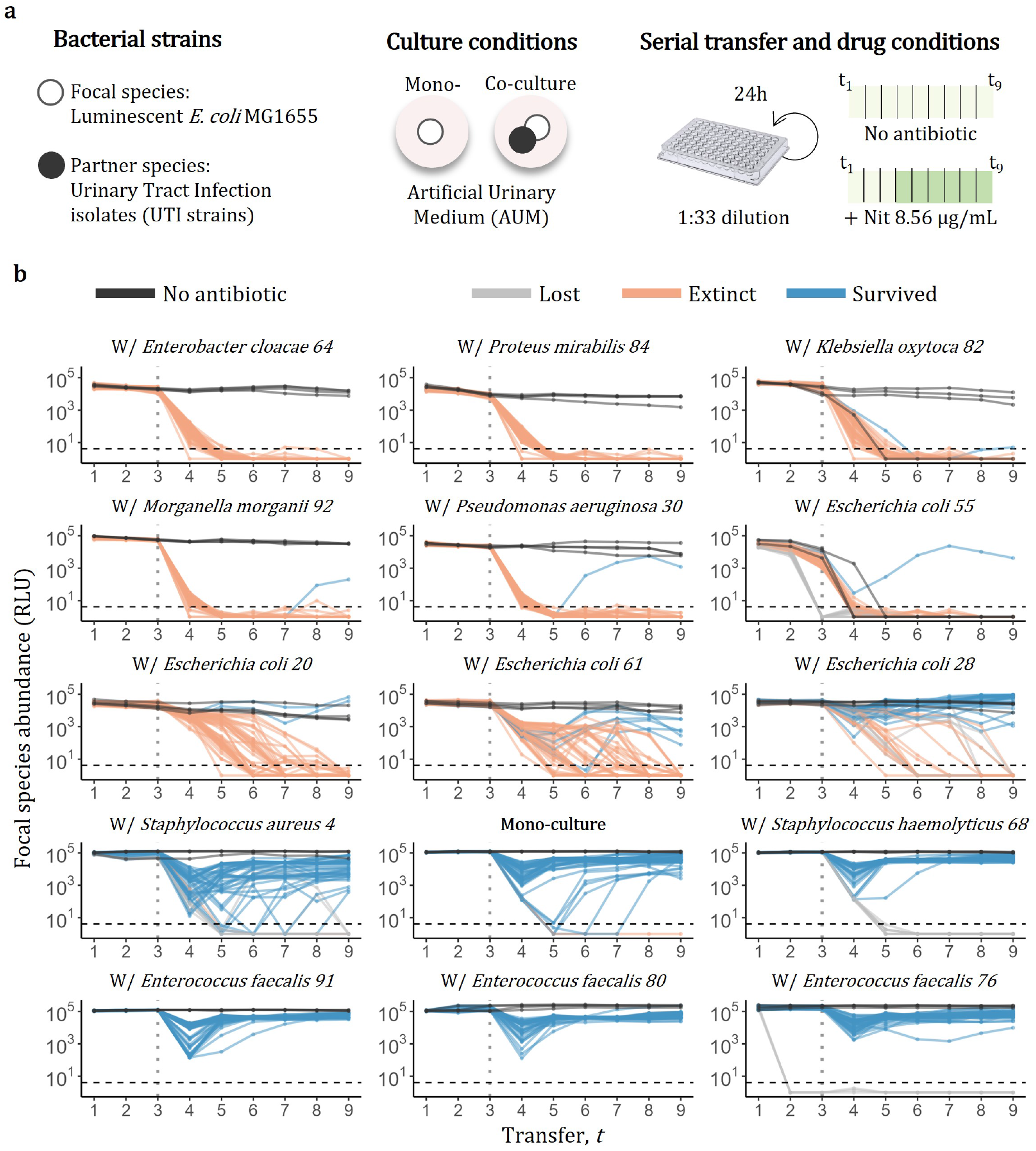
Evolutionary rescue under the influence of UTI isolates. **a)** Experimental setup. **b)** Dynamics of the focal strain (*E*.*coli* MG1655) measured via luminescence (RLU, relative light units). Partner species are indicated at the top of each plot. The horizontal dashed line corresponds to the limit of detection, while the vertical dashed line corresponds to the last time point without antibiotic. We considered “lost” those populations that went extinct before antibiotic exposure and those that were likely caused by experimental artifacts (in grey, *Methods*). We considered “extinct” those populations that were below the detection limit at the last transfer (in orange) and “survived” all the remaining ones (in blue). In black we show the populations (*n* = 4) that did not experience antibiotic.

In the absence of antibiotic, the focal *E. coli* strain was maintained at high abundance over the 9 transfers in all cases, except in co-culture with a second *E. coli* strain (isolate 55), where competition likely caused extinction independently of antibiotic use (black lines in Fig. **2**b). Upon antibiotic exposure, the focal *E. coli* declined in all conditions; some populations went below the detection limit (orange lines in Fig. **2**b), while others managed to survive and recover (blue lines in Fig. **2**b).

To test whether the observed recovery was driven by evolution of resistance, we characterized the growth phenotypes of evolved populations. Those that evolved under the antibiotic stress could grow to significantly higher yields at large nitrofurantoin concentrations compared to the focal strain evolved without antibiotic (Fig. S2) and to the ancestor (Fig. S3), confirming resistance gain. We consider the fraction of survived populations as a proxy for the probability of rescue and resistance evolution.

The presence of UTI isolates drastically affected the dynamics of the focal *E. coli* strain. When alone, it survived in 98% of the replicates. When together with one of the UTI isolates, the fraction of rescued populations covered the entire range from complete extinctions (e.g., in co-culture with *Enterobacter cloacae*) to complete rescue with all tested *Enterococcus faecalis* strains (Fig. **2**b, S4). The observed variation in the rescue dynamics highlights how strongly ecological interactions can affect the response to environmental challenges.

To test whether our model can explain and predict the observed variation in rescue probability, we inferred the partner-specific interaction parameters (*α* and *δ*), and then mapped each strain pair into the competition-protection space (*Methods* and Fig. **3**a-b). As predicted by our model, the fraction of observed rescue is greater or equal to the mono-culture in co-culture with the three *Enterococcus faecalis* isolates, which are the only ones with overall positive interactions (*α >* 1 −*δ*) mapping in the yellow region (Fig. **3**b). For all the other co-cultures, which had overall negative interactions (*α <* 1 −*δ*) mapping in the blue region, the fraction of observed rescue is smaller or equal to the mono-culture (Fig. **3**b and S4).

**Fig. 3:**
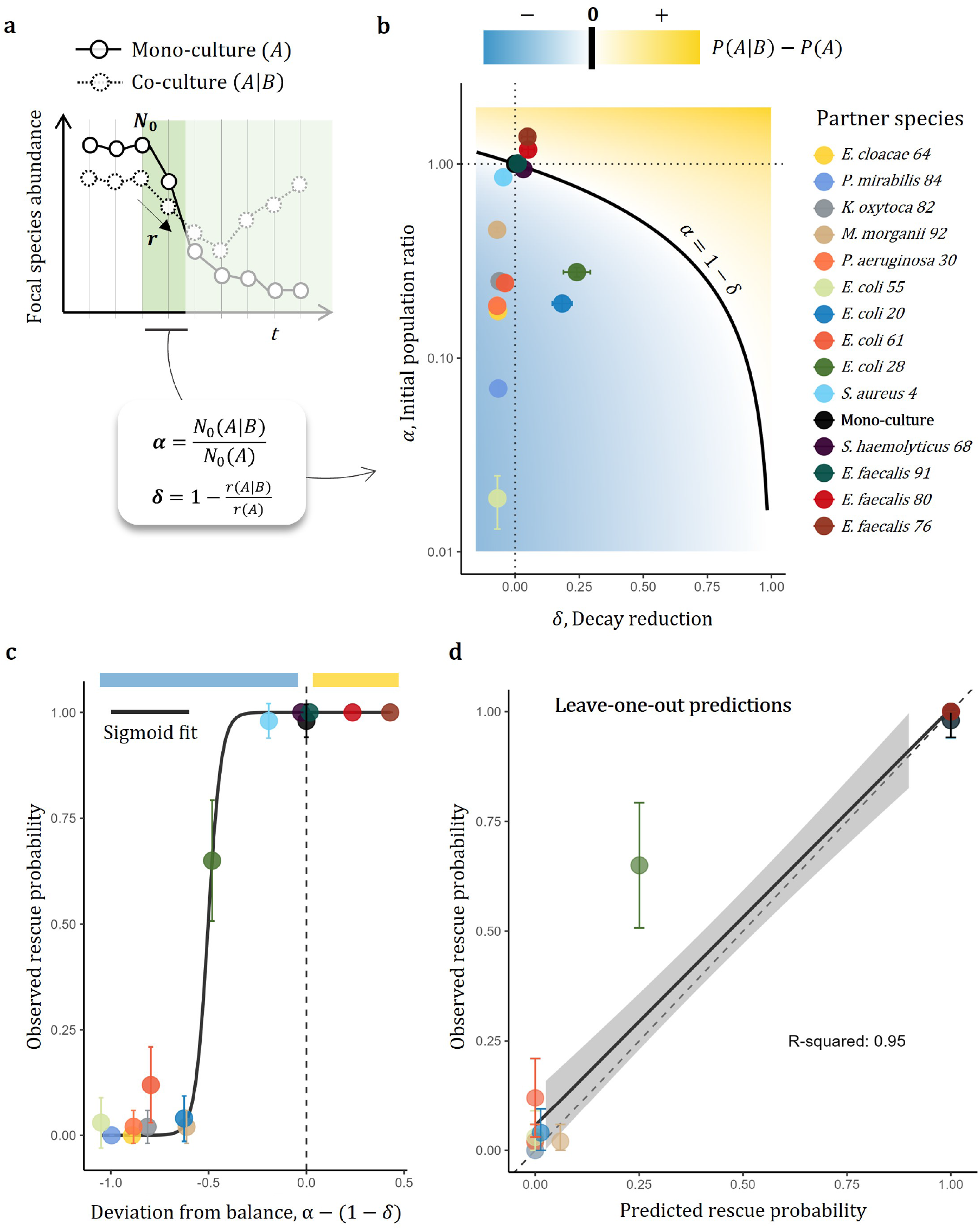
Evolutionary rescue can explain and predict the observed dynamics under the influence of UTI isolates. **a)** Schematics of how we infer the *α* and *δ* parameters from the dynamics (*Methods* for more details). **b)** Mapping of the UTI isolates into our competition-protection regimes. Model predict that the partner strains mapped in the blue region should hinder rescue of the focal strain, and vice versa for those in the yellow region. The solid black line represents the condition where the probability is expected to be equal in mono- and co-culture (i.e. the competition-protection balance) **c)** Correlation between the deviation from the competition-protection balance and the observed rescue probability. The solid line shows the sigmoid fit (bound between 0 and 1) used for the correlation, which fitted the data points with sum of squares due to regression, SSR=0.02. The vertical dashed line capture all strains that land on the solid black line of panel b. **d)** Leave-one-out predictions of rescue probability based on the sigmoid fit with the *α* and *δ* parameters after removing one of the species pairs, repeated for all pairs.

In addition to this qualitative prediction, we also expect that the weaker the ecological effect of a partner strain is, the closer the rescue probability should be to the mono-culture case. Indeed, the deviation from the competition-protection balance (*α* − (1 − *δ*)) strongly correlates with the observed rescue probability (Fig. **3**c), demonstrating that the two param-eters in our evolutionary rescue model can explain the observed variation in the rescue probability.

To test our model’s ability to predict the outcome of “unseen” co-cultures, we performed leave-one-out predictions: For each of the partner strains, we removed the corresponding data point and fitted a sigmoid curve to the remaining *n*− 1 strains to predict its rescue probability. This approach reveals that the model can almost perfectly predict the rescue probability of the focal strain in co-culture with a second out-of-sample strain, performing better at the extremes (where *P* is close to zero or one), and worse for intermediate values, likely due to the lack of data points with intermediate rescue probability (25 ≤*P* ≤75%) (Fig. **3**d).

Overall, the evolution of resistance to nitrofurantoin in the presence of UTI isolates can be explained by their effects on the focal strain’s initial population size and decay rate, as predicted by our minimal model of eco-evolutionary rescue. However, most of the variation, and explanatory power, comes from the effect on population size (*α*, Fig. **3**b), suggesting that competition, more than protection, is the main driver of AMR within the tested species pairs.

### Detoxification overcomes competition and shapes the evolution of resistance to ampicillin

To explore the potential and generality of the framework across unrelated bacterial ecosystems, we performed a second set of evolutionary experiments in a more controlled setting, where we could better explore the interplay between competitive and protective interactions, by manipulating the *α* and *δ* parameters through the medium composition.

We evolved two environmental strains (*Microbacterium liquefaciens Ml*, and *Comamonas testosteroni Ct*) in minimal medium with glucose and citrate, with or without the beta-lactam antibiotic ampi-cillin. Based on previous work [Piccardi et al., 2019, DosSantos et al., 2022], we expect that *Ct* and *Ml* will preferentially consume citrate and glucose respectively, limiting competition and ensuring coexistence via niche separation. In addition, *Ct* can detoxify ampicillin by producing beta-lactamase enzymes, which should make the environment more tolerable for *Ml* (Fig. **4**a).

**Fig. 4:**
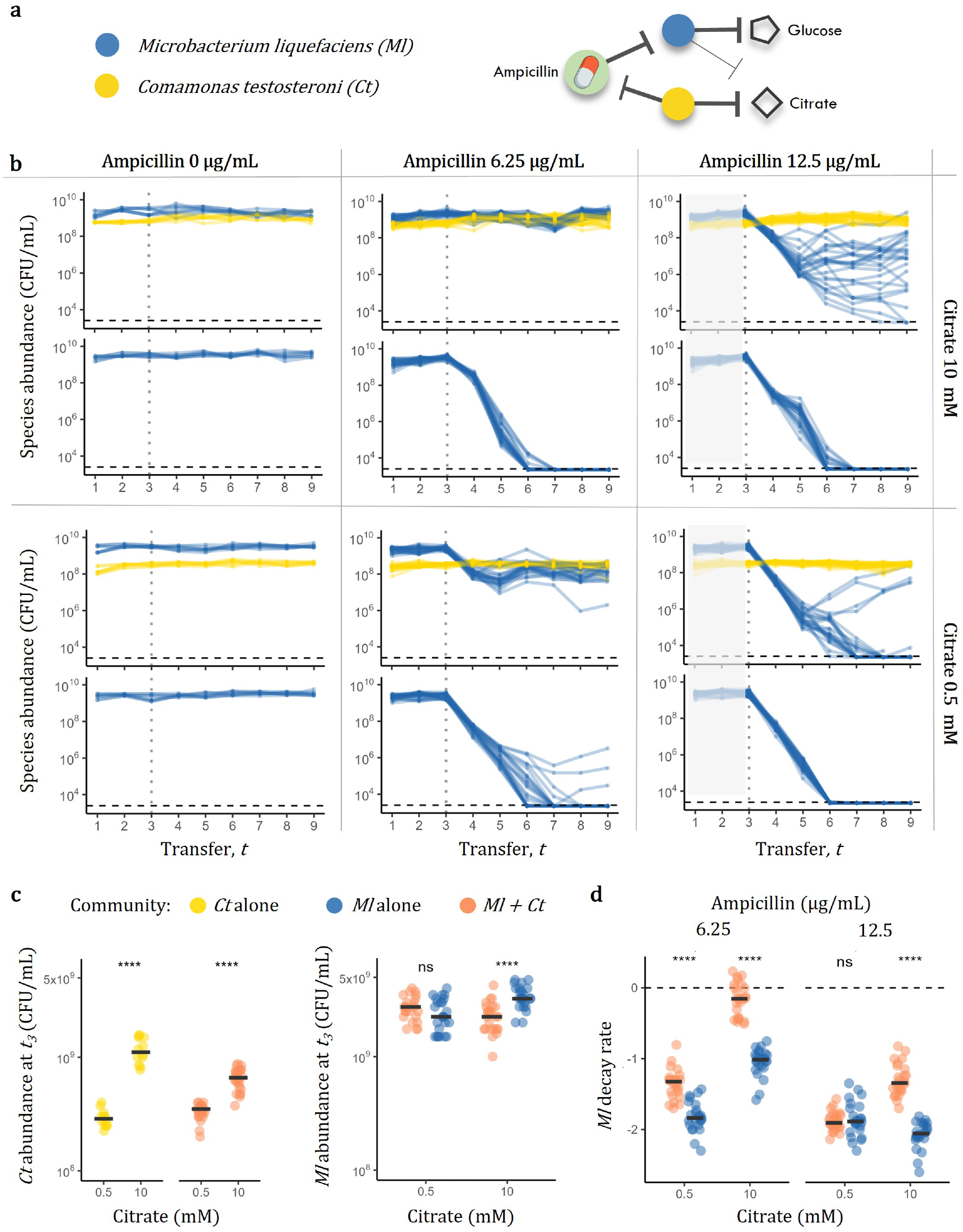
Dynamics under competition and detoxification. **a)** Schematics of species-resource-antibiotic interactions. **b)** Dynamics of *Ml* (in blue) and *Ct* (in gold) measured via colony forming units (CFU/mL) at the end of each transfer across the different conditions: increasing ampicillin concentration from left to right and increasing citrate concentration from bottom to top. The horizontal dashed lines show the limit of detection, while the vertical dotted lines show to the last time point without antibiotic. The gray areas in the ampicillin 12.5 condition highlight the fact that those data points are the same of the ampicillin 6.25 condition. This is because populations were split for the 2 antibiotic conditions only after the third transfer. **c)** Abundance of *Ct* and *Ml* at the last transfer before antibiotic exposure (*t*_3_), evolved either alone or together as specified by the colors. **d)** Decay rate of *Ml* (computed as the slope of the *log*_10_ dynamics after exposure to ampicillin). Colors indicate whether the transfers were performed in isolation (blue) or in co-culture with *Ct* (orange). The dashed line shows the no-decay slope, while the solid black lines show the condition-specific medians. Asterisks show significance level after t-test.

In mono-culture, the main drivers of extinction are the dilution factor and the antibiotic concentration, but populations may recover through evolutionary rescue (Fig. S5). In co-culture, the detoxification efficacy of *Ct* should depend on antibiotic as well as citrate concentration: higher ampicillin concentrations should take longer to be cleared, while higher citrate concentrations should speed up the detoxification process by enabling *Ct* to grow to larger populations. To test this and to compare rescue proba-bilities in mono- and co-culture, we serially passaged 24 independent replicates of *Ml* evolving alone or together with *Ct* in a minimal medium containing 10 or 0.5 mM of citrate (+ 15mM of glucose), every 24 hours over 9 dilution-growth cycles with dilution factor 1:50. The first three passages were antibiotic-free and the remaining six either continued without antibiotic, as a control, or were supplemented with 6.25 or 12.5 μg/mL of ampicillin (Fig. **4**b).

First, focusing on the dynamics prior to antibiotic exposure, we confirmed that lowering the citrate concentration (from 10 to 0.5 mM) decreases the steadystate abundance of *Ct* (Fig. **4**c, left), which in turn reduces the competition with *Ml* (Fig. **4**c, right) and should limit detoxification. After adding ampicillin, the presence of *Ct* reduces the decay rate of *Ml* under high citrate, while under low citrate the reduction is weaker or not significant (Fig. **4**d). All together, the data demonstrate how ampicillin and citrate concentrations modulate both the competitive and protective effects of *Ct*, which we hypothesized to be key for the evolutionary rescue dynamics.

After calculating the ecological effects of *Ct* through the *α* and *δ* parameters (Table **3**), our evolutionary rescue model predicts that the presence of *Ct* should (i) favor rescue in the two conditions with low ampicillin (6.25 μg/mL) (where *α >* 1 −*δ*), (ii) should not impact rescue when ampicillin was high (12.5 μg/mL) and citrate low (0.5 mM) (where both decay and initial population size of *Ml* do not change significantly, *α* ∼1 −*δ*), and (iii) should hinder rescue when ampicillin and citrate were high (12.5 μg/mL and 10 mM, respectively) (where *α <* 1 −*δ*). This is not what we observe: Upon antibiotic exposure, *Ml* extinctions in mono-culture always outnumber those in co-culture with *Ct*, regardless of the tested antibiotic or citrate conditions (Fig. **4**b and Fig. S6). This suggests that the overall effects of detoxification overcome competition, and that our minimal model lacks components explaining the observed dynamics.

### Strong or weak detoxification respectively lead to ecological or evolutionary rescue

We reasoned that two non-mutually exclusive processes could be taking place: First, *Ct* might ensure the survival of *Ml* populations independently of genetic changes through ecological processes alone (ecological rescue). Second, if rescue occurs due to genetic changes (evolutionary rescue), detoxification might affect not only the decay rate of the susceptible population, but also the selection acting on such resistant variants.

How can detoxification affect selection? To gain a quantitative answer to this question, we built a mechanistic model of growth by resource consumption, death by antibiotic and detoxification (*Methods*).

We simulated growth-and-dilution cycles mimicking the experiment and quantified how the presence of a detoxifying species can change the invasion success of emerging mutants. Simulations demonstrate that partially resistant mutants can get lost in monocultures, but invade in co-cultures with a detoxifying species, even when detoxification is strong enough to ensure survival of the susceptible type (Fig. **5**a).

**Fig. 5:**
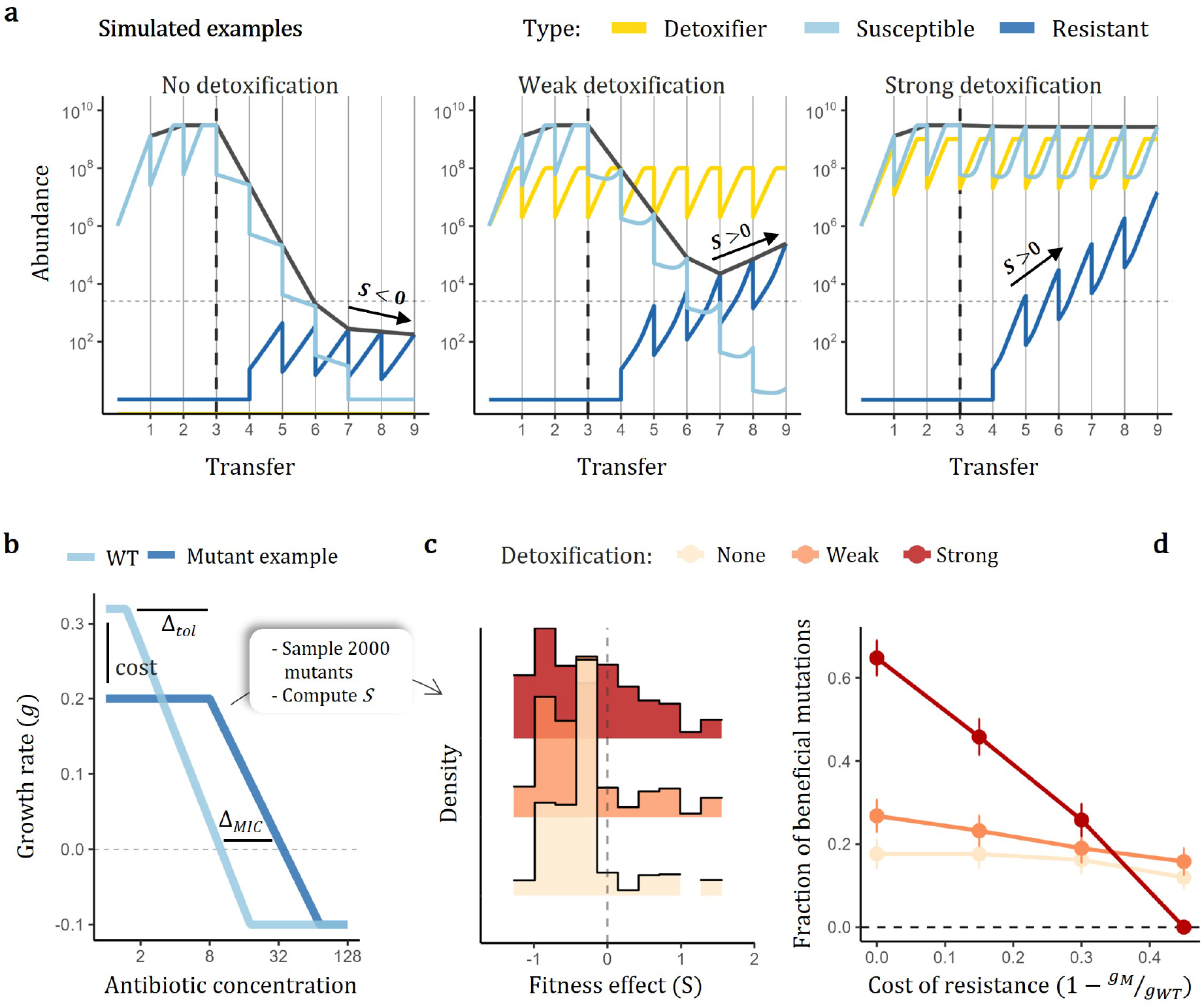
A mechanistic model of competition, detoxification and resistance to quantify the effect of detoxification on selection. **a)** Simulated examples of population dynamics across transfers, whereby the same resistant mutant does not invade in mono-culture but does invade under weak or strong detoxification. **b)** Response curves for a susceptible wild type (WT) and a resistant type (Mutant) defined by 3 parameters: the *cost* represents the decrease of growth in the absence of antibiotics (*cost* = 1 −*g*_*M*_ */g*_*W T*_), *tol* represents the maximum antibiotic concentration that does not affect growth and *MIC* represents the minimum concentration that prevents growth. After fixing the response curve of the WT, we simulate 2000 mutants by varying the cost value from 0 to 0.45, and sampling the remaining two parameters at random, more specifically: Δ*tol* ∼*Norm*(0, 10), Δ*MIC* ∼*Norm*(0, 15), and then compute the fitness effect of such mutations as the slope across serial transfers (*Methods*). **c)** Distribution of fitness under no, weak, or strong detoxification. **d)** Fraction of beneficial mutations (*S >* 0) as a function of the underlying cost and the detoxification strength. For these simulations we fixed: [Ab]=12.5 μg/mL as in the experiment, [Citrate]=0,1,10 to manipulate the strength of detoxification. (*Methods* for the full list of parameters value).

To generalize this finding, we simulated invasion dynamics for thousands of different mutants and quantified the strength of selection (measured as the slope between transfers) in the presence or absence of the detoxifying species. Assuming that each mutant has its own growth response to the antibiotic, which we defined as a function of cost, tolerance and minimum inhibitory concentration (Fig. **5**b, *Methods*), our simulations show that the fraction of mutations that can successfully invade and rescue the population (*S >* 0) increases with detoxification (Fig. **5**c). Importantly, mutations that carry a strong cost are instead less likely to invade under strong detoxification, as this increases the time during which the cost matters, i.e. the absence of antibiotic (Fig. **5**c). This result suggests that detoxification can enlarge the pool of resistant mutants, by making the environment tolerable to weakly resistant types, which carry a small cost, supporting our hypothesis. For our evolutionary rescue framework, this means that we cannot ignore the effect that the partner species has on the input of beneficial mutations (the *β* parameter, Fig. **1**b).

To test whether the observed rescue dynamics are due to the evolution of ampicillin resistance, we isolated the *Ml* populations that survived the 9 transfers with antibiotic and quantified their growth in ampicillin. Overall, the response curves vary depending on whether *Ml* evolved alone or in co-culture, with low or high citrate, and with low or high ampicillin (Fig. **6**a). The *Ml* populations that evolved with 0.5 mM citrate significantly outperform the ancestral population when growing under the antibiotic conditions that match the evolution experiments (ampicillin 6.25 or 12.5 μg/mL), while the *Ml* populations that evolved with 10 mM citrate do not (Fig. **6**b). These data show how weak detoxification can facilitate the evolution of resistance, while strong detoxification can promote survival without resistance, through *ecological* rescue alone.

**Fig. 6:**
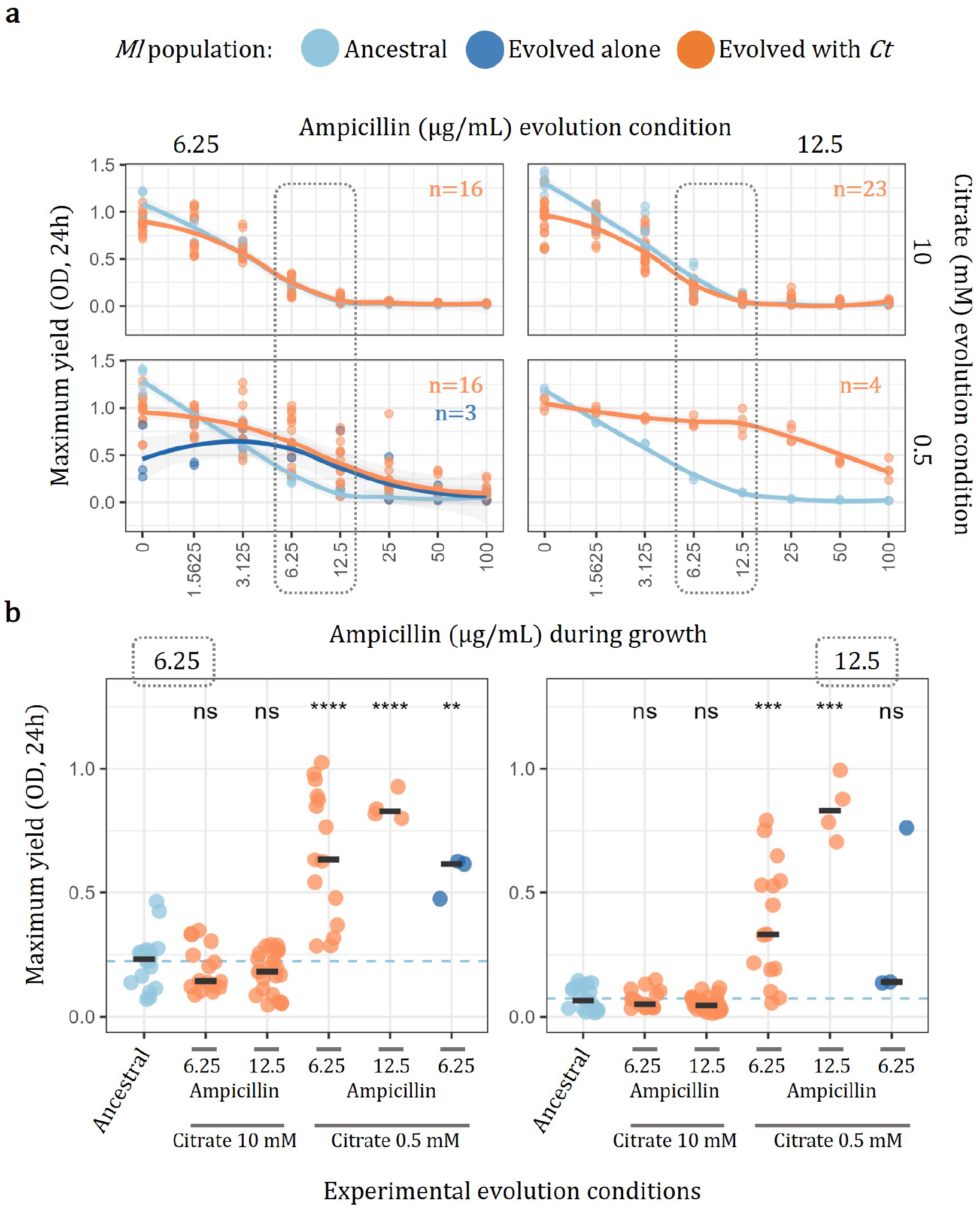
Ampicillin susceptibility of evolved *Ml* populations. **a)** Maximum population size of *Ml*, calculated as max achieved optical density (OD) within 24 hour growth on minimal medium + glucose (15mM). The different quadrants organize the populations based on the conditions of their transfer experiment: Ampi-cillin 6.25 or 12.5 μg/mL from left to right and citrate 0.5 or 10 mM from bottom to top. Number of tested evolved populations is indicated for each condition (different numbers are due to the availability of populations which survived the transfers). Each conditions contains 2 replicates of the control (ancestral *Ml*) grown in the same plate. Solid lines represent the smoothed conditional mean. **b)** Zoom on the maximum yield of *Ml* upon exposure to the ampicillin concentrations that were used during our evolution experiments (6.25 or 12.5 μg/mL). Here, the different replicates of ancestral *Ml* were pulled together. The dashed line shows the mean of the ancestral populations, while the solid black lines show the condition-specific medians. Asterisks show significance level after t-test comparison against the control.

As a proof of principle, our mechanistic consumer-resource model with detoxification can recapitulate the observed dynamics at high citrate and high ampicillin, whereby the susceptible population first drops and then slowly recovers, without any evolutionary process, but simply assuming a gradual accumulation of the enzyme responsible for detoxifica-tion (Fig. S7, *Supplementary materials*).

Finally, putting all the data together and mapping them to our evolutionary rescue framework, we show that the effects on initial population size and decay rate alone fail to explain why more survival is observed in the strong-antibiotic-strong-detoxification regime (red triangle in Fig. **7**a). When we add the effect that detoxification by *Ct* can have on the distribution of mutation effects of *Ml*, the model correctly predicts increased evolutionary rescue in all co-culture regimes (Fig. **7**b), but still fails to explain the lack of increased resistance in the populations evolved under strong detoxification (Fig. **6**b). Only when we assume that strong detoxification causes ecological rescue (i.e. *δ* = 1 such that the susceptible type’s decay is reduced to zero) does our framework explain the experimental dynamics (Fig. **7**c), whereby the presence of a detoxifying species always promotes the survival of the susceptible species, but not necessarily via genetic changes.

**Fig. 7:**
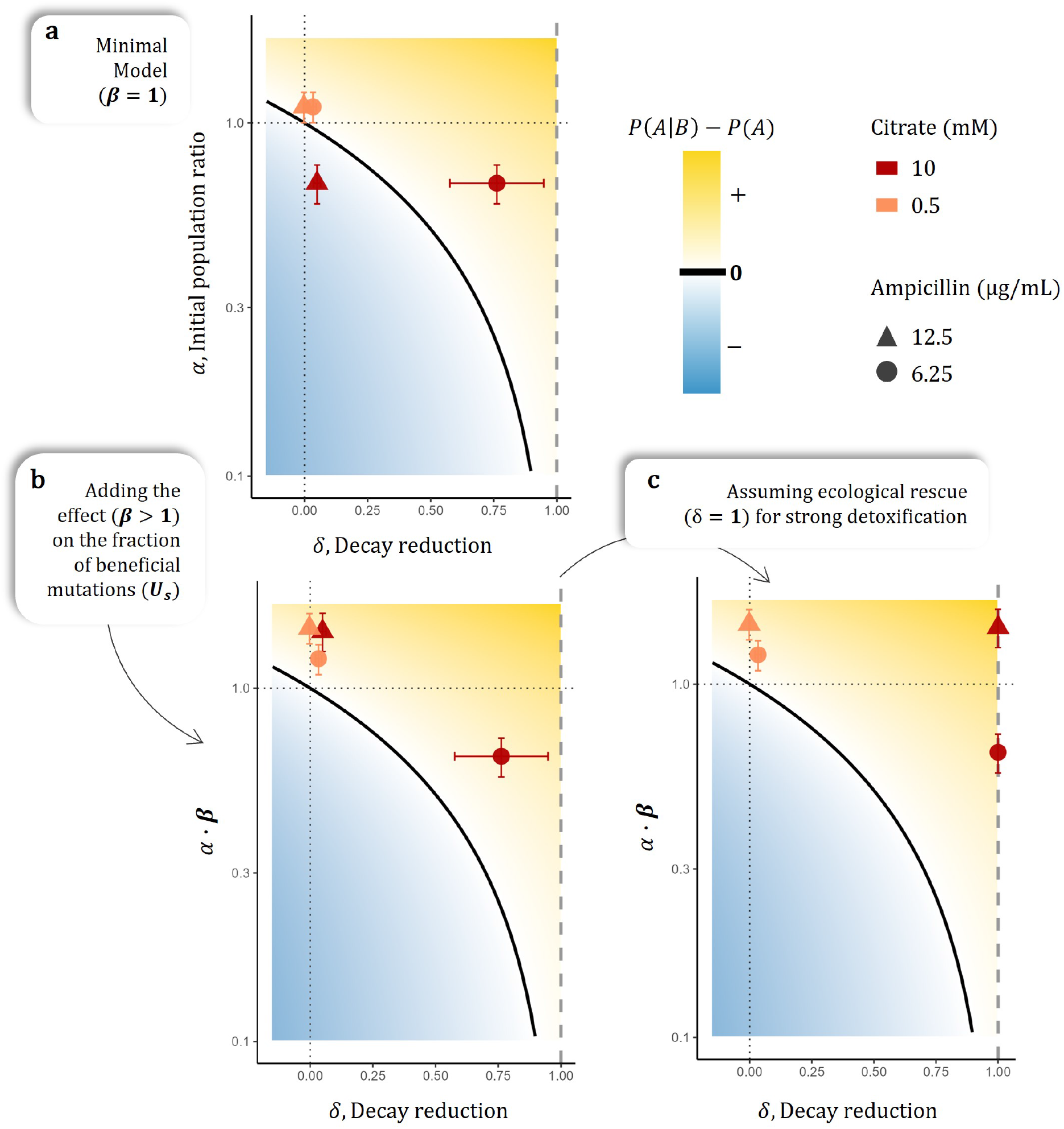
Competition and protection effects on rescue probability. **a)** Mapping of the effects of *Ct* on the initial population size of *Ml* and on its decay rate (*α* and *δ*). **b)** Mapping of the effects of *Ct*, including the expected increase in the fraction of beneficial mutations (*β*). **c)** Mapping of the effects of *Ct*, assuming that the strong detoxification regime (10 mM citrate) led to ecological rescues (*δ* = 1). Data points represent the mean computed across 24 independent replicates per condition, and the bars represent the standard error around the mean (calculated from the experimental data for *α* and *δ*). The symbols color and shape identify different experimental conditions (citrate and ampicillin concentrations, respectively). The solid black line represent the condition *α · β* = 1 *− δ* (i.e. the competition-protection balance).

## Discussion

Our work provides a quantitative framework to compare the evolution of resistance in isolation or in the presence of interacting species. First, expanding classical evolutionary rescue theory, we identified the conditions under which ecological interactions can favor the emergence of resistance. And second, through experimental evolution of different bacterial species, we validated our model’s predictions, demonstrating how the balance between competition and protection controls the likelihood that microbial populations can resist exposure to antibiotics.

To first validate our theoretical results, we studied the evolution of resistance to nitrofurantoin in *E. coli* MG1655 in the presence of different bacterial strains isolated from patients with urinary tract infections (UTIs). Our experiments suggest that competitive effects between UTI isolates are key drivers of AMR evolution. All the UTI isolates tested reduced the survival probability of the focal *E. coli*, with the exceptions of *Enterococci* and *Staphylococci*, consistent with previous measurements of interactions among UTI strains [DeVos et al., 2017]. A recent study similarly showed how the rate of antibiotic resistance evolution in *E. coli* increases in the spent medium of *Enterococcus faecium* and *Staphylococcus haemolyticus*, but decreases in the spent medium of a *Klebsiella* species [Zandbergen et al., 2024], again in agreement with our observations (Fig. **2**-**3**). Future work could investigate the specific mechanisms underlying both interactions and resistance, and extend the study to different bacterial families.

While the UTI isolates helped to validate our model in a clinically relevant system, we were unable to explore competitive and protective interactions simultaneously. A second set of experiments with a pair of environmental bacteria showed how protection by a beta-lactamase-producing species (*C. testosteroni, Ct*) always overcame competition for resources, improving the survival of the susceptible species (*M. liquefaciens, Ml*) in all co-culture conditions compared to the mono-cultures.

The observation of a susceptible species being protected by others is not new. Several studies show how beta-lactamase-producing bacteria can promote the survival of otherwise susceptible species (e.g. [Perlin et al., 2009, Bottery et al., 2022]). What is new, is how we can quantitatively integrate this process in the context of multiple ecological interactions and the resistance probability. For example, under the guidance of the model, we tuned the citrate concentration to manipulate the competitive and protective effects of *Ct*, which in turn shaped *Ml* ‘s response to ampicillin, showing how weak detoxification favored survival through the acquisition of resistance, while strong detoxification favored survival independently of resistance evolution, but through ecological rescue (i.e. the complete removal of the stressor).

To describe the evolution of resistance in our simple microbial communities, we made use of two different mathematical approaches. The first one, based on evolutionary rescue theory, relies on a few parameters to capture dynamics that could generalize across different natural and experimental settings [Orr and Unckless, 2008], while the second one, adapting classical consumer-resource models [Macarthur, 1970], describes explicit mechanisms of interactions and growth (such as detoxification and serial transfers) to capture dynamics that are specific to our experimental system. The latter complemented the minimal model of evolutionary rescue, but its results do not necessarily generalize to other systems. For example, our mechanistic model suggests that, even upon ecological rescue, antibiotic resistance might still emerge due to the transient but repeated exposure to antibiotics (Fig. **5**). This holds true for dynamics with serial re-introduction of antibiotics (such as growth-and-dilution experiments) but might differ for dynamics where detoxification brings antibiotic concentration to a constant low steady state (such as chemostat experiments).

While our main goal was to learn how ecological interactions shape the relative likelihood of evolving resistance (i.e. co-culture relative to mono-culture), we focused less on absolute probabilities. The conditions tested with our experiments represent a window where ecological interactions have an effect. However, in more extreme scenarios (e.g. when the probability to acquire resistance is very high), the effects of ecological interactions might be harder to detect (simulated examples are shown in Fig. S8). To predict the absolute probability that a population evolves resistance remains a difficult task as it requires computing non-trivial quantities, such as the rate of spontaneous mutations and the distribution of their effects, both of which might depend on specific environmental conditions.

Applications of evolutionary rescue theory span from natural to experimental populations, and from conservation biology to medicine [Alexander et al., 2014]. These dynamics have been studied through different mathematical formulations (e.g. [Gomulkiewicz and Holt, 1995, Lasky, 2019, Azevedo and Olofsson, 2021]), and under a variety of different assumptions (such as population structure [Aif et al., 2022], multiple mutations [Martin et al., 2013] or time-dependent stresses [Marrec and Bitbol, 2020]). Here, we expand on the seminal work by Orr and Unckless, which relies on several simplifying assumptions, but provides easy-to-interpret and easy-to-test results [Orr and Unckless, 2008, Orr and Unckless, 2014]. Importantly, we assume that resistance arises through a single mutation of fixed effect, which grows independently from co-occurring genotypes, in a well-mixed environment, with non-overlapping generations (i.e. discrete dynamics), and under a constant stress. The explicit form of the rescue probability further relies on the assumption of weak selection (*s, r* ≪1) [Orr and Unckless, 2008], which is most likely not met under strong antibiotic regimes. This could lead to an overestimation of rescue probabilities [Azevedo and Olofsson, 2021]. However, as discussed above, because we are interested in relative probabilities, this discrepancy should cancel out when comparing mono- with co-cultures.

To account for ecological effects, we further assume that the partner species’ dynamics do not depend on the stress, nor on the other species. Taking into account these additional factors gives rise to density-dependent and/or co-evolutionary dynamics whereby the ecological interactions are not constant in time (theoretical [Osmond and DeMazancourt, 2012, Elzen et al., 2017] and experimental examples [Melero-Jiménez et al., 2025]). We briefly explore such scenarios (*Supplementary materials*), but recognize that more work is needed to fully characterize these out-of-equilibrium dynamics, especially for communities with more than two species (e.g. [DeMazancourt et al., 2008, Low-Décarie et al., 2015]).

Besides providing quantitative predictions, our model guided us to identify key factors controlling the evolution of resistance in the presence of interacting species. In fact, we argue that the main strength of our framework is not its predictive power, but rather its exploratory support: When the observations do not match the predictions, one is forced to explore previously overlooked aspects, and in doing so, we gained new knowledge about our system of interest.

In our case, the first minimal formulation of the evolutionary rescue model, based on the ecological effects on population size (*α*) and the decay rate (*δ*) alone, could describe the evolutionary response of *E. coli* to nitrofurantoin, but was not enough to fully explain how *Ml* could survive ampicillin under competition and detoxification by *Ct*. This led us to hypothesize that detoxification might affect resistance evolution in additional ways: by favoring the invasion of weak resistant mutations and/or by fully removing the source of stress (i.e. ecological rescue). Simulating serial transfer dynamics, we compared the distributions of fitness effects (DFE) under different ecological regimes and confirmed that detoxification can indeed increase the fraction of beneficial mutations. We do not argue that the simulated DFEs represent the “real” ones, but they served as proof-of-principle for how even a small increase in the input of beneficial mutations could favor rescue in co-culture (Fig. **7**b). This result is in line with the recurring observation that low antibiotic concentrations favor resistance evolution [Ramsayer et al., 2013, Lindsey et al., 2013], only in our case the lowering of the concentration is achieved by a partner species.

In this work, we mainly focused on ecological interactions controlling competition and protection, but depending on the biological system, it will be important to integrate additional effects into the evolutionary rescue framework. For example, when comparing AMR evolution in mono- or in co-cultures with populations carrying resistance on plasmids, or other mobile elements, it is necessary to include horizontal gene transfer (HGT) [Frost et al., 2005]. One way this can be done in our framework is to consider HGT as an additional source of resistant mutations and quantify this effect through the parameter *β*. In the extreme cases with high HGT rates (*β*≫ 1), cocultures may favor rescue and the spread of AMR, regardless of competition.

Despite our focus on AMR dynamics, the proposed framework should similarly hold for different sources of stress (e.g. temperature, salinity or pH) [Melero-Jiménez et al., 2025], but may need to be adjusted depending on the relevant factors. For example, under temperature increase, protective effects might be less relevant compared to competition or selection as it is unlikely that the presence of a second species mitigates the stress imposed by higher temperature [Toll-Riera et al., 2022] (but see [Frank, 2020]).

To conclude, combining phenomenological models of evolutionary rescue with mechanistic models of ecological interactions and experimental evolution with bacteria, we describe how competitive and protective interactions control the evolution of resistance within simple microbial communities. Our work provides simple and quantitative rules to guide future empirical and theoretical studies to further investigate the evolution of resistance to antibiotics –and other environmental challenges – in the context of species-rich ecosystems.

## Methods

### Evolutionary rescue theory

Here, we extend a classical evolutionary rescue framework [Orr and Unckless, 2008] to include ecological interactions and compare the probability that a focal species *A* will survive in isolation or in the presence of other species. For the main results we focus on a 2-species ecosystem, and assume *B* to be a single partner species. We assume that the focal species population is composed of two types: susceptible (*a*) or resistant (*a*′) to the environmental stress. Upon stress, the most common genotype in the population *a* decays geometrically: *N*_*t*_(*a*) = *N*_*t−*1_(*a*)(1 −*r*), with *r >* 0. Instead, the resistant genotype *a*′ grows in the stressful condition as: *N*_*t*_(*a*′) = *N*_*t−*1_(*a*′)(1 + *s*), with *s >* 0. Importantly, we assume the growth of the mutant to be independent from the decay of the wild type, contrary to the original formulation by Orr and Unckless whereby the mutant grows by *N*_*t*_(*a*′) = *N*_*t−*1_(*a*′)(1− *r* + *S*) [Orr and Unckless, 2008, Orr and Unckless, 2014]. While the dynamical systems are equivalent (rewriting *s* = *S*− *r*), the choice of the parameter definition influence the interpretation of the ecological effects on them. We decided to model the absolute growth of the resistant genotype *a*′ independently from the decay rate *r* to avoid not well justified correlations. In Fig. S10 we show how the shape of the competition-protection balance changes under different assumptions on the relationship between *r* and *s*. Our evolutionary rescue framework includes drift as sampling noise (individuals of genotype *a*′ at time *t* are drawn from a Poisson distribution with mean *N*_*t−*1_(*a*′) · (1 + *s*)). Importantly, the resistant variant *a*′ can either be already present in the population prior to stress exposure at a given frequency *p*_0_, as part of the standing genetic variation (SGV), or emerge after stress exposure via *de novo* mutation (dnM). We consider the probability of rescue via both standing genetic variation (*P*_sgv_) and via *de novo* mutation (*P*_dnm_) and compare such probabilities in isolation (*P* (*A*)) or in the presence of a partner species (*P* (*A* |*B*)).

The rescue probabilities of our focal species *A*, in the absence of ecological interactions, can be approximated by the following:

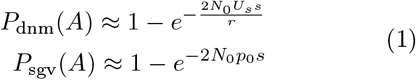

The key parameters are therefore: the decay rate (*r*), the beneficial effect of the mutation (*s*), the rate of *de novo* mutations with such an effect (*U*_*s*_), the initial number of individuals (*N*_0_) and the frequency of resistant mutants (*p*_0_) when the environment changes (*t* = 0).

For the rest of the study we assume that neither *A* nor the environmental stress affect the partner species *B*. In the *Supplementary materials* we study how breaking this assumption leads to dynamical systems where the effects on the rescue probabilities are density-dependent. We define *α, β, γ, δ, ϵ* as the constant effects of *B* on *N*_0_, *U*_*s*_, *p*_0_, *r* and *s*, respectively (definitions in Tables **1**-**2**). The probability of observing evolutionary rescue in the presence of ecological interactions with *B* becomes:

**Table 1:** Species, genotypes and variables definition.

| Name | Interpretation |
| --- | --- |
| $A$ | Focal species |
| $B$ | Partner species |
| $a$ | Susceptible genotype of species $A$ |
| $a'$ | Resistant genotype of species $A$ |
| $W_i$ | Fitness of genotype $i$ |
| $s$ | Growth advantage of $a'$ |
| $r$ | Decay rate of $a$ |
| $N_t$ | Total population size of $A$ at time $t$ |
| $U_s$ | Rate of mutations with effect $s$ |
| $p_0$ | Proportion of resistant types at time 0 |
| $c$ | Cost of resistant mutation prior to stress |

**Table 2:**
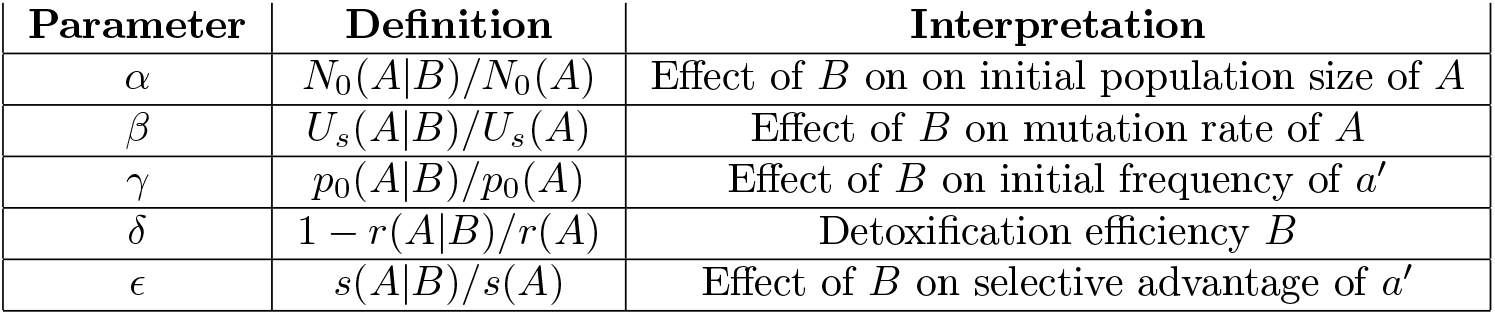
Definition of the ecological effects of species *B* on species *A*.

**Table 3:**
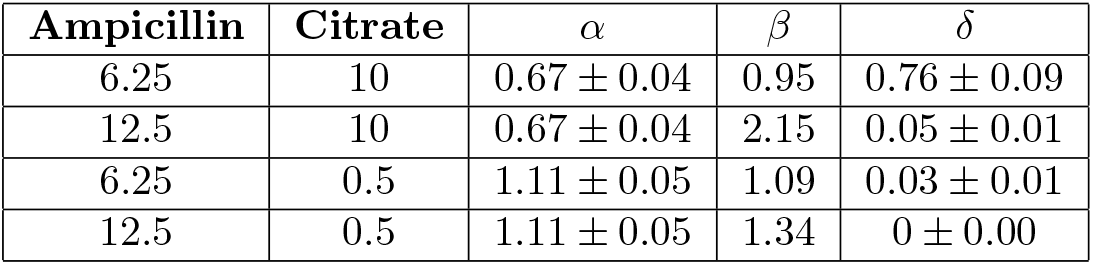
Inferred interaction parameters.

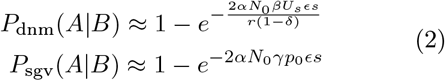

Comparing the **Eq.1** and **2**, it follows that:

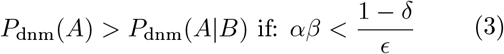

and for the rescue via standing genetic variation:

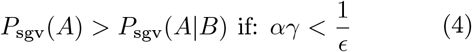

The total rescue probability is given by a combination of rescue via *de novo* mutations and standing genetic variation: *P*_tot_ = *P*_sgv_ +(1 −*P*_sgv_)*P*_dnm_. Now combining the two from **Eq.1 and 2**, we get:

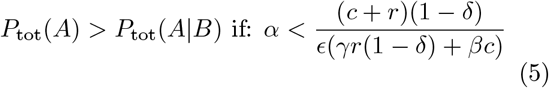

where *c* represents the relative contribution of *de novo* and existing mutational input (*c* = *U*_*s*_*/p*_0_). Assuming that the partner species does not affect selection strength, nor the supply of beneficial mutations (*β, γ, ϵ* = 1), we obtained the competition-protection balance described in the Results section and shown in Fig. **1**.

### Mapping experiments on the competition-protection space

We quantified the previously defined interactions *α* and *δ* as follows: For each partner species *i* under experimental condition *j* in replicate *k*, the ecological effect on the population size of the focal species *A* is 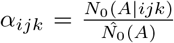. Here *N*_0_(*A*|*ijk*) is the observed population size of *A* for that specific experimental replicate before antibiotic exposure (time *t*_3_), while 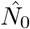 (*A*) is the mean population size of *A* in mono-cultures before antibiotic exposure (time *t*_3_). Then, *α*_*ij*_ is the mean competition effect across replicates 1 to *k*. Similarly, the ecological effect on the decay rate is 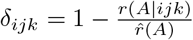, where *r*(*A*|*ijk*) is the observed decay rate *A* for that specific experimental replicate, quantified as 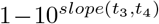. The exponential transformation is because we compute the slope on the *log*_10_ of the abundance *N*. Then, 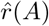is the mean decay rate of *A* in mono-cultures and *δ*_*ij*_ is the mean protective effect of partner *i* under condition *j* across replicates 1 to *k*.

### Mechanistic model of growth by resource consumption, detoxification and resistance

Here, we build a mechanistic model that can reproduce our experimental setup in order to study the effects of a detoxifying partner species on the decay rate and the selection acting on a focal species. For this purpose we expanded classical consumer-resource models [Macarthur, 1970] by considering also the dynamics of toxins (i.e. antibiotics) which can kill a species and/or be detoxified by another. More specifically, we model Monod dynamics of two species (*A* and *B*), two resource types (*R*_1_ and *R*_2_) and of one toxin (*T*). The growth dynamics follow serial batch cultures of 24 hours with a 1:50 dilution, as in our experiment. Assuming that species *B* is resistant to the range of toxin concentrations under study and consumes *R*_2_ only, while species *A* is susceptible to the range of toxin concentrations under study and consumes *R*_1_ only, the dynamics take the following form:

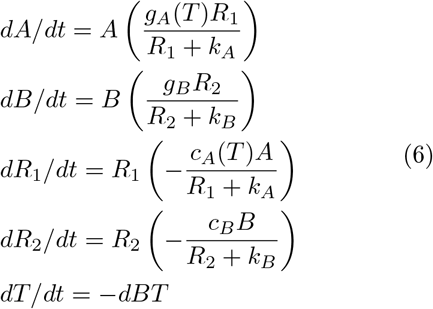

Here, *dX/dt* represents the change of *X* over an infinitesimally short time, *g*_*X*_ (*T*) describes the max growth rate of species *X* as a function of the toxin concentration *T, k*_*X*_ represents the half saturation constant of species *X, c*_*X*_ represents the consumption rate by species *X*, and finally *d* represents the detoxification rate of species *B*. The consumption rate of a resource is related to the growth rate as follows: *g*_*X*_ = *c*_*X*_*Y*_*X*_, where *Y*_*X*_ represents the yield (resource to biomass transformation coefficient). We assume that the toxin *T* affects the growth rate of species *A* through a step-like response curve. In particular, the step function is defined by 3 parameters: the max growth rate achievable (*g*_*max*_), the minimum toxin concentration at which the growth is affected (hereafter *tol*) and the minimum toxin concentration at which growth is positive (hereafter *MIC*). So the growth rate of species *A* takes the following shape:

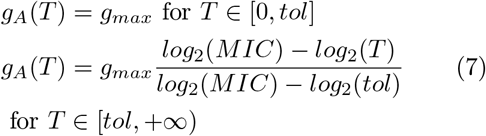

According to our definition of the response curve, growth is not affected for any toxin concentration *T < tol*, and decays exponentially for any toxin concentration *T > tol*. In particular it is equal to 0 or negative for *T* ≥*MIC*. This acts as a step with a linear decrease when taking the *log*_2_ of toxin values (Fig. **5**b). Importantly, given our model definition, we assume that antibiotic affects growth through the yield, and not consumption rate.

The parameters used to produce the simulations for Fig. **5**a are as follows: *g*_*W T*_ = 0.32, *g*_*mut*_ = 0.3, *g*_*B*_ =0.27, *Y*_*W T*_ = *Y*_*mut*_ = 2 ·10^8^, *Y*_*B*_ = 10^8^, *d* = 10^*−*9^, *T* =12.5,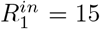, 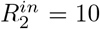, *k*_*A*_ = 1, *k*_*B*_ = 0.1, *tol*_*W T*_ =1.5, *tol*_*mut*_ = 7.5, *MIC*_*W T*_ = 10, *MIC*_*mut*_ = 25, where the wild type *WT* and the mutant *mut* represent two sub-populations of species *A*.

### Quantifying selection over random mutants

Making use of the mechanistic model presented above, we generate thousands of mutants, simulate their growth under serial batch cycles and quantify their invasion selection. Each mutant is defined by its phenotype *g*_*mut*_ = *g*_*W T*_ (1 −*cost*), *tol*_*mut*_ =*tol*_*W T*_ + Δ_*tol*_ and *MIC*_*mut*_ = *MIC*_*W T*_ + Δ_*MIC*_, whereby the cost is 0, 0.15, 0.3 or 0.45, and the Δ are drawn from zero-centered Gaussian distributions:Δ_*tol*_ ∼ *N*(*μ* = 0, *σ*= 10), Δ_*MIC*_ ∼ *N*(*μ* = 0, *σ*= 15). With these assumptions we simulate mono- or co-cultures where *T* = 0 for the first 3 transfers, then *T* = 6.25 or 12.5 and the input of the resource consumed by the detoxifying species (i.e. citrate)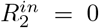, 1 or 10, to simulate different degrees of detoxification. We assume that the mutant emerges at transfer 4 and study its dynamics afterwards. We measure selection acting on each mutant as the slope of its abundance over transfers, and not within transfers. This is because even if a mutant grows within a transfer it does not ensure survival over transfers due to dilution. Repeating this procedure for 2000 randomly generated mutants (500 per each cost value), we quantify the distribution of fitness effects with no, weak or strong detoxification as in Fig. **5**c, and quantify the fraction of beneficial mutations (Fig. **5**d).

## Experiments with UTI partners

### Strain collection and growth media

The focal *Escherichia coli* strain is a derivative of MG1655 with a chromosomal integration of the *luxCDABE* operon, introduced using plasmid pBJ6 (Addgene plasmid 167135) as described by [Matsumoto et al., 2022]. This focal strain was grown in mono-culture or co-culture with bacterial strains from a previously characterized collection of 72 isolates from patients with polymicrobial urinary tract infection (UTI) [Croxall et al., 2011, DeVos et al., 2017]. Strains were grown in artificial urine medium (AUM; [Brooks and Keevil, 1997]) with or without nitrofurantoin (Sigma N7878).

### UTI isolates selection criteria

Out of the 72 available UTI isolates, we chose 14 to balance the number of replicates (50 per partner strain) and the experimental feasibility based on previous characterization (Espinosa et al. in preparation). To select these 14 isolates we prioritize three aspects: they should be at least partially resistant to the chosen nitrofurantoin concentration (8.56 μg/mL); they should display potentially different interactions with the focal strain; and should span across different taxonomic groups. After excluding the UTI isolates whose yield was reduced more than half by nitrofurantoin 10.4 μg/mL, we finally chose all UTI isolates with positive effects on the focal strain’s yield and/or or response to nitrofurantoin (*Enterococcus faecalis 76, 80, 91, Escherichia coli 55, 20, 61, 28, Staphylococcus haemolyticus 68, Klebsiella oxyloca 82*) and the remaining 5 isolates having mild or strong negative interactions on both yield and response to antibiotic (*Staphylococcus aureus 4, Morganella morganii 92, Proteus mirabilis 84, Pseudomonas aeruginosa 30, Enterobacter cloacae 64*).

### Evolutionary rescue setup and population dynamics read-out

Glycerol stocks from single colonies of each selected UTI isolate were distributed in a 96-well microtiter. A complementary plate was prepared with glycerol stock of the focal strain. The plates design consisted of an array of randomly distributed wells in such a way that when mixed, in 6 replicates per plate of blank wells, the focal strain in mono-culture or in co-culture with each isolate were obtained. Before the experiment, overnight cultures were initiated by inoculating both 96-well plates independently and incubating for 48 hours to assure saturation. Throughout the experiment, plates consisted of 200 μL per well and incubation was done at 30°;C with constant shaking. The first transfer was performed by pinning the saturated overnights in a 1:1 ratio into nine 96-well plates with fresh AUM to a 1:1000 dilution. Subsequent transfers were carried out every 24 hours, transferring 6 μL of culture into fresh medium. The first two transfers were performed in fresh AUM; subsequent transfers were made in either AUM (4 replicates per condition) or AUM supplemented with nitrofurantoin (NIT) at a final concentration of 8.56 μg/mL (50 replicates per condition). An experimental plate was frozen at −80°;C with a final concentration of 15% glycerol. The growth of the focal strain was monitored by measuring luminescence every 30 minutes after vigorous shaking. Luminescence crosstalk between wells was corrected following the approach described by [Kishony and Leibler, 2003]. Population yield was quantified by calculating the median of the four highest luminescence readings per culture after each transfer. The detection limit to classify extinction was defined as the maximum observed RLU (relative light units) among the control wells (∼5 RLU, corresponding to ∼20 colony formation units per well). Extinction and survival of focal and UTI strains were further confirmed by plating from last transfer on agar (CHROMagar Orientation, CHROMagar; results not shown).

### Automatic platform artifact identification and removal

Transfers were performed by a robotic automatized platform. Errors in the pipetting device (e.g. by tip clogs or channel malfunction) happen and result in systematic loss of cultures. A first visual examination of the data indicated that such artifacts occurred during the experiment, as populations were lost from otherwise stable cultures in particular plate positions. To identify artifacts with a consistent methodical strategy, we compared the number of observed extinctions in every well with the number we would expect by chance. If we observed more extinctions than expected occurring in the same well position, an artifact was flagged. More precisely, for each community *i* (mono-culture of the focal strain + 14 pairs with the UTI partners) we count the number of extinctions (*x*_*i*_) and assume *P*_*i*_ = *x*_*i*_*/N*, where *N* = 50 is the number of replicates, to be the “true” extinction probability of *i*. Then, for each well (*wi*) containing community *i*, we count the number of extinctions (*x*_*wi*_) and quantify the probability of observing at least *x*_*wi*_ events given *P*_*i*_. This conditional probability of observing at least *x* events in *n* trials can be computed as: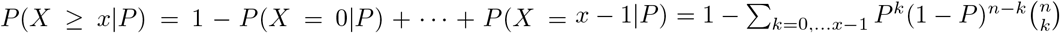, where in our case *n* = 9 as the number of 96-well plates used. If such a probability is small (we arbitrarily set *P* (*X*≥ *x*_*wi*_ |*P*_*i*_) *<* 0.05), then observing *x*_*wi*_ extinctions is rare enough that we consider it a potential artifact and we remove the extinctions from that well from the analysis. This test resulted in 4 wells (out of 96) showing signals of potential artifacts. The grey lines in Fig. **2** show the replicates that were removed from subsequent analysis.

### Resistance gain characterization after evolution

First, a simple characterization to distinguish between populations evolved under no-drug and drug treatment consisted in growth of three experimental plates from the last transfer day (4 and 14 replicates of no-drug and drug treatments) by pinning into fresh AUM or AUM with 17 *μ*g/ml nitrofurantoin in microtiter plates. Plates were incubated for 48 hours at 30°;C with constant shaking, measuring luminescence every 40 minutes after vigorous shaking. Maximal luminescence (median of the highest three luminescence reads for each well) was compared among treatments and populations. Second, dose response curves were done using one plate (6 replicates for each culture condition) from the last experimental day as described above, but pinning into plates with the different concentrations of antibiotic. Dose response curves (DRC) for each culture were obtained by normalizing growth to the yield in AUM without drugs.

### Rescue probability correlation

To test whether our model could explain the rescue probability across communities, we correlated the observed fraction of survived populations (*P*) with combinations of the key parameters identified by our model: *α* and *δ*. Because probabilities by definition are contained in the [0, 1] range, we performed the correlation by fitting a sigmoid curve bound between 0 and 1, which takes the following shape: 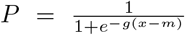, where *g* corresponds to the growth rate, *m* corresponds to the mid-point of the sigmoidal and *x* is our predictive variable. In particular, we show that the combined variable *x* = *α* − (1− *δ*) accurately describes the variation in the observed probabilities. Importantly, *x* represents the deviation from the null-effect line, in fact, when *x* = 0 rescue in mono-culture should equal rescue in co-culture, when *x <* 0 mono-cultures should show more rescue and vice versa when *x >* 0.

### Leave-one-out predictions

To test whether our model could predict the rescue probability within “unseen” species pairs, we performed a leave-one-out approach (shown in Fig. **3**d) as follows: for each community *i*, we removed the corresponding probability of rescue from the data set, then we used the remaining 14 communities to fit a new sigmoidal curve as explained above and as shown in Fig. **3**c, and finally we used this fitted curve to predict the rescue probability within community *i*, given the deviation from the null-effect line caused by *i* (*α*_*i*_ − (1 − *δ*_*i*_)). We then report the correlation (and the corresponding *R*^2^) between the predicted and the observed rescue probabilities.

### Experiments with detoxifying partner

#### Strain collection and growth conditions

For this set of experiments we used *Microbacterium liquefaciens Ml* and *Comamonas testosteroni Ct*, two bacterial species previously isolated from industrial settings and characterized in laboratory conditions [Piccardi et al., 2019, DosSantos et al., 2022]. *Ml* lacks the ability to synthesize thiamine, biotin, L-proline and L-cysteine, thus we supplement them in all the minimal media used in our experiments (respectively at concentration 0.5 μg/mL, 0.1 μg/mL, 0.15 mg/mL and 0.15 mg/mL) [Sulheim et al., 2026]. The medium used throughout the experiment consisted of these 4 supplemented amino acids and vitamins + HMB, M9 (see [Martino et al., 2024] for recipes) + glucose (10 mM) and citrate (10 or 0.5 mM). Before starting the evolution experiment, we isolated colonies on TSA plates, cultured the two species individually in 10mL of minimal medium supplemented with glucose and 10 mM of citrate for two rounds of 48 hours. This step was performed to obtain pre-adapted populations whose growth on minimal medium is reproducible, which we use to seed all the presented serial transfer experiments (i.e. the ancestral populations at *t*_0_).

#### Evolutionary rescue setup and population dynamics read-out

Our evolutionary rescue setup consisted of two main serial transfer experiments (one with 10 and one with 0.5 mM of citrate), where we performed 9 growth cycles of 24 hours followed by 1:50 dilution. More specifically, we grew our pre-adapted populations of *Ml* and *Ct* on TSA plates, picked multiple colonies and let them grow in 10 mL of H_2_O with 4%TSB, for 24 hours under shaking conditions (200RPM) at 28*°;*C. After this step, we inoculated 4μL of bacterial population into each well of a 96-well plate, each containing 196μL of the specific medium condition, to start the transfer experiment. For each transfer we passaged 4μL from the old into the newly prepared 96-well plate and grew the populations at 28*°;*C, shaking incubator (200RPM) for 24 hours. At the end of each transfer, we plated several dilutions of each well on selective media to count colony forming units (CFUs): we counted *Ml* on TSA + Colistin (10 μg/mL) plates, which prevent the growth of *Ct*, and we counted *Ct* on TSA plates, where *Ml* does not form visible colonies within 24 hours. We quantified the population sizes of *Ml* and *Ct* over 9 transfers, across different media and antibiotic conditions. For each medium condition (10 or 0.5 mM of citrate + 15mM of glucose), we grew all populations without antibiotics during the first 3 growth cycles, then from transfer 4 to 9 we added either 6.25 or 12.5 μg/mL of ampicillin. For each medium-antibiotic combination, we had 10 populations evolving without ampi-cillin (4 *Ml* mono-cultures, 4 *Ml-Ct* co-cultures and 2 *Ct* mono-cultures) and 60 evolving with ampicillin (24 *Ml* mono-cultures, 24 *Ml-Ct* co-cultures and 12 *Ct* mono-cultures) for a total of 280 communities (70*×* 2 citrate concentrations *×*2 ampicillin concentrations) over 9 time points. The two antibiotic concentrations were tested in the same experiment by duplicating the 96-well plate from transfer 4 onward. For graphical convenience, the data related to the first 3 transfers of the same medium condition were duplicated (shadowed area in Fig. **4**b).

### Response curves after evolution

To evaluate susceptibility of the evolve populations of *Ml*, we measured the growth performance of each survived population and compared them against the ancestral (pre-adapted *Ml* at *t*_0_). We plated the frozen populations on TSA + Colistin (10 µg/mL) plates, which should exclude *Ct*, pre-cultured them for 24 hours in 10mL of H_2_O with 4%TSB, under shaking conditions (200RPM) at 28°C. To control for contamination by *Ct* we plated the populations on TSA plates and discarded the replicates where *Ct* colonies could be observed. We inoculated 4µL of bacterial culture in each well of 96-well plates containing minimal medium + glucose 15mM and a gradient of ampicillin ranging from 100 to 0 µg/mL by a factor of 2, and grew all of them under shaking conditions at 28°C to measure their optical density (OD_600_) every 10 minutes for 48 hours. To test for an increase in susceptibility we compared the maximum achieved OD within the first 24 hours (equivalent to the evolutionary transfer duration) of all the evolved populations against the ancestral.

## Data availability

Data and scripts are available on GitHub at https://github.com/Mitri-lab/resistance-competition-protection.

## Competing interests

Authors declare that they have no competing interests.

## Acknowledgments

We thank Afra Salazar, Eric Ulrich, Margaret Vogel, Alessia Del Panta, Prajwal Padmanabha, Snorre Sulheim and Alan Pacheco for feedback on earlier versions of the manuscript. AEC, GP, and TB were supported in part by German Research Foundation (DFG) Collaborative Research Centre (SFB) 1310 and DFG standalone grant BO 3502/4-1 (project number 498531688). MA and SM were funded by the Swiss National Science Foundation Eccellenza Grant PCEGP3_181272, project grant 236360 and the NCCR Microbiomes SNF 51NF40_180575.

## Authors contribution

**MA** designed the overall project; built the mathematical models; performed the experiments related to the evolution of resistance to ampicillin; analyzed both experimental and simulations data; produced figures and statistical analysis; wrote the first draft of the manuscript and edited the following versions. **AEC** co-designed, performed, and co-analyzed the experiments with the UTI-collection. Reviewed and edited the manuscript. **GP** provided assistance with the robotic platform (UTI experiments). **TB** supervised (UTI experiments); acquired funding; reviewed and edited the manuscript. **SM** supervised the overall project; acquired funding; wrote, reviewed and edited the manuscript.

## Supplementary materials

### Parameters definition

#### Evolutionary rescue theory and its extension

##### Introducing the evolutionary rescue model

Classic evolutionary rescue theories study the fate of a population that upon an environmental challenge risks extinction, but the emergence of genetically resistant variants can revert the process and rescue the population. Here, we extend such a framework to include ecological interactions and compare the probability that a focal species *A* will survive in isolation (*P* (*A*)) or in the presence of other species (*P* (*A* |*B*)). For the main results we focus on a 2-species ecosystem, and assume *B* to be a single partner species. Following [Orr and Unckless, 2008], we assume that the focal species population is composed of two types: susceptible (*a*) or resistant (*a*′) to the environmental stress. Upon stress, the most common genotype in the population (hereafter *a*) decays geometrically: *N*_*t*_(*a*) = *N*_*t−*1_(*a*)(1 *−r*), where *r >* 0 is the decay rate due to the stress. In contrast, the resistant genotype *a*′ grows in the stressful condition as: *N*_*t*_(*a*′) = *N*_*t−*1_(*a*′)(1 + *s*), assuming the selective advantage *s >* 0. Importantly, we assume the growth of the mutant to be independent from the decay of the wild type, contrary to the original formulation whereby the mutant grows by *N*_*t*_(*a*′) = *N*_*t−*1_(*a*′)(1 −*r* + *S*). While the dynamical systems are equivalent (re-writing *s* = *S −r*), the choice of the parameter definition influence the interpretation of the ecological effects on them. Evolutionary rescue is commonly characterized by a U-shaped curve representing the total abundance of the endangered population (**Fig. 1a**). The uncontrolled growth of the mutant is unrealistic, however the probability of rescue, which is the main focus of our work, is equivalent to the probability that a population reaches a reasonable *N*\*, large enough to escape drift. As far as the population grows above *N*\*, what happens after does not influence the rescue probabilities and is beyond the scopes of this work. In the absence of demographic noise (drift), any genotype with *s >* 0 would establish and rescue the population. However, because the resistant mutations appear in few individuals, variation can be lost by chance even if beneficial. Our evolutionary rescue framework includes drift as sampling noise (individuals of genotype *a*′ at time *t* are assumed Poisson-distributed with mean *N*_*t−*1_(*a*′)(1 + *s*))). Importantly, the resistant variant *a*′ can either be already present in the population prior to stress exposure at a given frequency *p*_0_, as part of the standing genetic variation (SGV), or emerge after stress exposure via *de novo* mutation (dnM). We consider the probability of rescue via both standing genetic variation (*P*_sgv_) and via *de novo* mutation (*P*_dnm_) and compare such probabilities in isolation (*P* (*A*)) or in the presence of a partner species (*P* (*A* |*B*)). The rescue probabilities of our focal species *A*, in the absence of ecological interactions, can be approximated by the following:

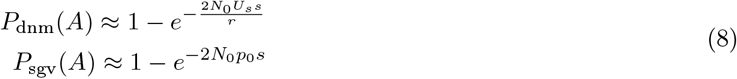

##### Adding ecological interactions

Next we determine the conditions under which rescue is more likely in isolation or in the presence of other species *B* (when *P* (*A*) *> P* (*A* |*B*)). For the rest of the study we assume that neither *A* nor the environmental stress affect the partner species *B*. Breaking this assumption leads to dynamical systems where the effects on the rescue probabilities are density-dependent (as discussed later on). We define *α, β, γ, δ, ϵ* as the constant effects of *B* on the initial inoculum of *A* (*N*_0_), the rate at which the resistant allele emerge *de novo* (*U*_*s*_), its frequency prior to the environmental change (*p*_0_), the decay rate of the susceptible genotype (*r*) and the selection benefit of the resistant one (*s*), respectively (definitions in Tables **1**-**2**). The rescue probabilities of our focal species *A* to experience evolutionary rescue in the presence of ecological interactions with *B* become:

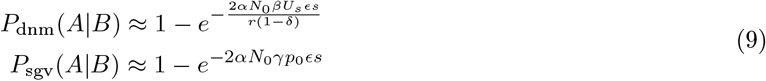

Comparing the **Eq.1** and **2**, it follows that:

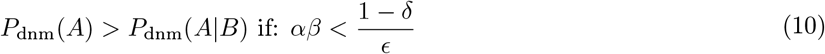

and for the rescue via standing genetic variation:

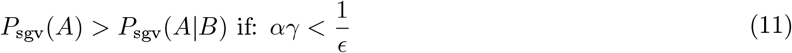

The total rescue probability is given by a combination of rescue via *de novo* mutations and standing genetic variation: *P*_tot_ = *P*_sgv_ + (1 *− P*_sgv_)*P*_dnm_. Now combining the two from **Eq.1 and 2**, we get:

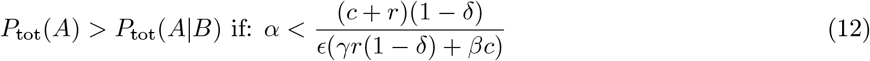

where *c* represents the relative contribution of *de novo* and existing mutational input (*c* = *U*_*s*_*/p*_0_).

##### The competition-protection balance

To narrow down the exploration of multiple parameters, we focused on the *α* and *δ* parameters, because those are the easiest to measure and control experimentally and because we hypothesized that they encompass the most common interaction types within species-rich communities. *α*, defined as the effect of *B* on the initial population of *A*, can capture all those interactions that affect growth prior to environmental stress: 0 *< α <* 1 correspond to competition (e.g. driven by niche overlap), while *α >* 1 correspond to facilitation (e.g. driven by niche creation). *δ*, defined as the effect of *B* on the decay rate of *A*, captures those interactions that can mitigate or exacerbate the environmental stress: if *δ <* 0 the environment is made harsher by the presence of *B*, while if 0 *< δ <* 1 the environment is made more tolerable. Notice that the condition *δ* = 1 represents the extreme case of complete detoxification, whereby the wild type population does not decay and the probability of rescue is not defined. Comparing the rescue probabilities in isolation or in the presence of *B*, we find that the relative strength of competition and protection determines whether resistance is favored (Competition-Protection balance):

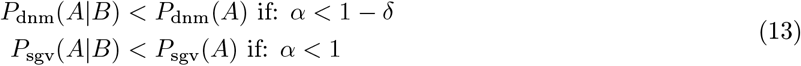

The curves in **Fig. 1c** represent the transitions from a parameter region where the evolution of resistance is more likely in isolation, to a region where this is more likely in the presence of a partner species. While the actual probabilities of rescue depend on all the parameters (*N*_0_, *U*_*s*_, *p*_0_, *r, s*, Fig. S8), these transitions only depend on the biotic interactions (*α, δ*) (Eq.13).

Combining the rescue probabilities as before, we obtain:

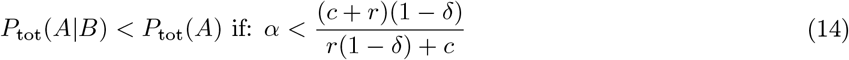

Here, the condition for which the rescue probability in isolation equals the rescue probability in co-culture depends on the *c* parameter too (the relative contribution of new mutations (*U*_*s*_) and standing variation *p*_0_).However, if there is detoxification (0 *< δ <* 1) it holds that: 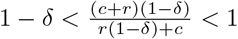 so that the transition curve must sit in between the the two defined in Eq.13. For small *c(c →* 0) the condition approaches the standing genetic variation case, while for large *c* the condition approaches the *de novo* mutation case (Fig. **1**c).

##### Mutation-selection balance and the cost of resistance

Under mutation-selection balance—the long-term equilibrium between mutational input (*U*_*s*_) and the strength of purifying selection (here *c*)—deleterious alleles can be maintained at low frequencies (here, *p*_0_ = *U*_*s*_*/c*). Because resistant mutations often carry a cost (a growth defect in the absence of the stress), the previously defined parameter *c* = *U*_*s*_*/p*_0_ can be interpreted as the cost associated with a deleterious mutation that would stabilize at frequency *p*_0_ under mutation-selection balance. Following this interpretation, costly resistant mutations are not expected to contribute to standing genetic variation due to strong purifying selection, but should rather emerge *de novo*. Recent studies have shown that the presence of additional bacterial species can modulate the cost of resistance in different ways [Cardoso et al., 2020, Klümper et al., 2019]. If the partner species affects the cost of having a resistant mutation in the absence of the stress by *ρ* = *c*(*A*|*B*)*/c*(*A*), then the initial proportion of resistant alleles is *p*_0_(*A*|*B*) = *p*_0_(*A*)*/ρ*, and the condition in **Eq.13** becomes:

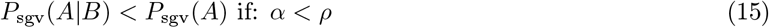

If the partner species *B* exacerbates the cost (*ρ >* 1), it makes it harder for resistance to establish from standing variation. If such exacerbation is strong enough, resistance can be hindered even if the partner species detoxifies the environment and facilitates the focal species. But, if facilitation (e.g. via cross-feeding) is strong enough (*α > ρ*), then *B* always promotes the rescue of *A*, regardless of detoxification. Intuitively, the larger the cost, the smaller is the contribution of standing variation to the combined rescue probability, and vice versa (**Fig. 1c**). Assuming mutation-selection balance might be reasonable for populations in the wild, which have evolved for long enough time under a given condition. However, populations studied in the lab are often generated after a single or few growth cycles of an initially isogenic population. This means that not enough time has passed for mutations to accumulate and for selection to act on them. In a population of size *N*_0_, grown from a single individual, we expect *U*_*s*_*log*(*N*_0_) mutations with effect *s*. This means that costly mutations can contribute to standing genetic variation and play a larger role in the lab where populations are large and far from reaching mutation-selection balance.

##### Density-dependent interactions

So far, we have studied the case where the effect of species *B* on the decay is constant. This can be interpreted as the case where species *B* does not change in abundance, thus detoxification is constant. What happens if the detoxification efficiency is function of the abundance of *B*, and *B* is not constant in time? Let’s define the instantaneous detoxification efficiency as: *d*_*e*_(*t*) := *d*(*N*_*B*_(*t*)*/K*_*B*_), where *K*_*B*_ is the carrying capacity that species *B* can reach in isolation. For *N*_*B*_(*t*) = *K*_*B*_ we retrieve the previous formalization. Let’s assume that the abundance of *B* depends on that of *A*, for example due to competition, and let’s define *α*_*ij*_ as the effect of species *i* on the carrying capacity of species *j*: *α*_*AB*_ = *K*_*B*|*A*_*/K*_*B*_ and equivalently *α*_*BA*_ = *K*_*A*|*B*_*/K*_*A*_. Notice that, assuming that at time *t*_0_ the populations are at steady state, this definition is equivalent to the previously defined interaction *α* = *N*_*A*|*B*_(*t*_0_)*/N*_*A*_(*t*_0_). Now we need to define a function *N*_*B*_(*t*) = *f* (*K*_*B*_, *N*_*A*_(*t*), *α*_*AB*_) such that:

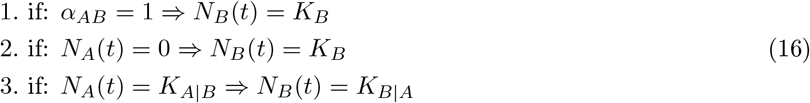

Let’s define the abundance of *B* as *N*_*B*_(*t*) := *K*_*B*_ *−N*_*A*_(*t*)*λ*_*AB*_. Assuming the the populations are at steady state upon environmental change (at *t*_0_), it follows that *N*_*A*_(*t*_0_) = *α*_*BA*_*K*_*A*_ = *K*_*A*|*B*_ and *N*_*B*_(*t*_0_) = *K*_*B*_ *− α*_*BA*_*K*_*A*_*λ*_*AB*_.

Then, to satisfy condition 3. from Eq.16, it follows that *K*_*B*_*− α*_*BA*_*K*_*A*_*λ*_*AB*_ = *α*_*AB*_*K*_*B*_, which solved for *λ* becomes:

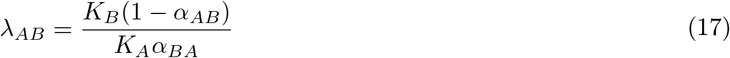

Thus, the dynamics of *B* become:

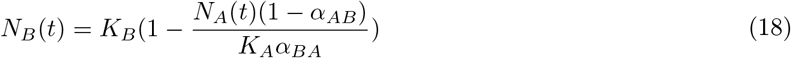

Such formulation satisfies all conditions of Eq.16 and can describe the dynamics of *B* as a function of *A* and the *α*_*ij*_ parameters. Now we can rewrite the dynamics of *A* including the density-dependent detoxification:

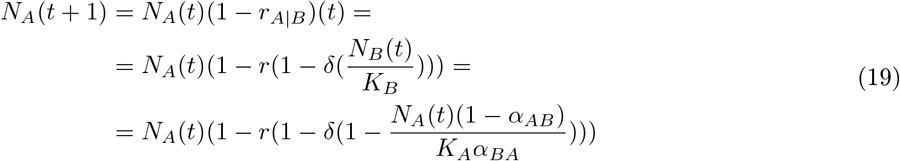

It follows that at time *t*_0_, *N*_*A*_ = *α*_*BA*_*K*_*A*_, so *N*_*A*_(1) = *N*_*A*_(0)(1− *r*(1− *δα*_*AB*_)) and if *A* competes with *B* (*α*_*AB*_ *<* 1), the *t*_0_ is the time point where the population of *B* is at its minimum and so is the effective detoxification. Thus, the effective density-dependent detoxification is bound between *δ· α*_*AB*_ and *δ*. In other words, if the population of the detoxifying species is reduced by the susceptible species, detoxification starts low at the beginning but increases over time as the susceptible species drops. Even though the probability of rescue depends on the full dynamics, we can use these boundary conditions to bound the probability too (Fig. S9).

### Mechanistic model of growth by resource consumption, detoxification and resistance

Here we build a mechanistic model that can reproduce our experimental setup in order to study the effects of a second detoxifying population (i.e. a partner species) on the distribution of fitness effects of mutations that emerge from the focal population (i.e. the focal species). For this purpose we expand classical consumer-resource models by considering also the dynamics of toxin (i.e. antibiotics) which can kill a species and/or be detoxified by another. More specifically, we model Monod dynamics of 2 species (*A* and *B*), 2 resource types (*R*_1_ and *R*_2_) and of 1 toxin (*T*). To mimic our experimental scenario, *A* is susceptible to the antibiotic, while *B* can degrade it. The growth dynamics follow serial batch cultures of 24 hours with a 1:50 dilution, as in our experiment. The dynamics take the following form:

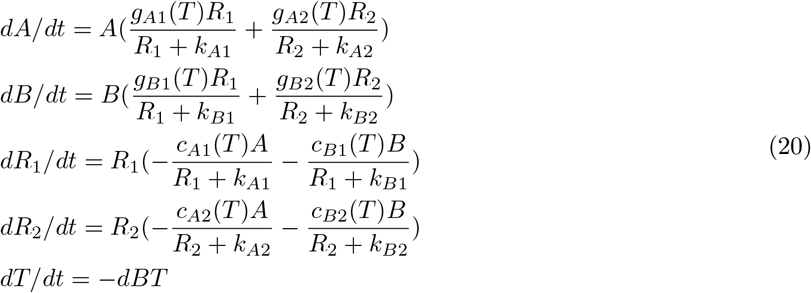

Here, *dX/dt* represents the change of *X* over an infinitesimally short time, *g*_*Xi*_(*T*) describes the max growth rate of species *X* on resource *i* as a function of the toxin concentration *T, k*_*Xi*_ represents the half saturation constant of species *X* on resource *i, c*_*Xi*_ represents the consumption rate of resource by species *x*, which is related to the growth rate by *g*_*Xi*_ = *y*_*Xi*_*· c*_*Xi*_, where *y*_*Xi*_ represents the yield (resource to biomass transformation) of species *X* on resource *i*, and finally *d* represents the detoxification rate of species *B*. We assume that the presence of a toxin *T* can affect the growth rate of a species and model this through a step-like response curve. In particular, the step function is defined by 3 parameters: the max growth rate achievable (*g*_*max*_), the minimum toxin concentration at which the growth is affected (hereafter tolerance: *tol*) and the minimum toxin concentration at which growth is positive (hereafter Minimum Inhibitory Concentration: *MIC*). So the growth rate of species

*A* takes the following shape:

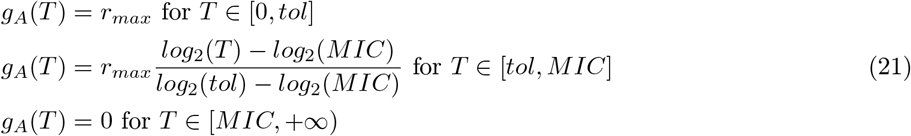

According to our definition of the response curve, growth is not affected for any toxin concentration below *tol*, it is equal to 0 for any toxin concentration above the *MIC*, and decays exponentially for any toxin concentration between *tol* and *MIC*. This looks as a step with a linear decrease when taking the *log*_2_ of toxin values (for example Fig. **5**b). For the rest of the work, we assume that species *B* is resistant to the range of toxin concentrations under study (so *g*_*B*_(*T*) = *g*_*B*_ *>* 0) and consumes *R*_2_ only, species *A* is susceptible to the range of toxin concentrations under study and consumes *R*_1_ only. We then study the fate of a mutant *M*, which generates from species *A* and enters the population at low abundance at a fixed point in time. We assume the mutant population *M* to have equal resource preferences as its wild type *A*, but it differs in the growth response to the toxin: *g*_*M*_ = *g*_*A*_ + Δ_*g*_, *tol*_*M*_ = *tol*_*A*_ + Δ_*t*_ and *MIC*_*M*_ = *MIC*_*A*_ + Δ_*m*_. Finally, the full dynamics are given by:

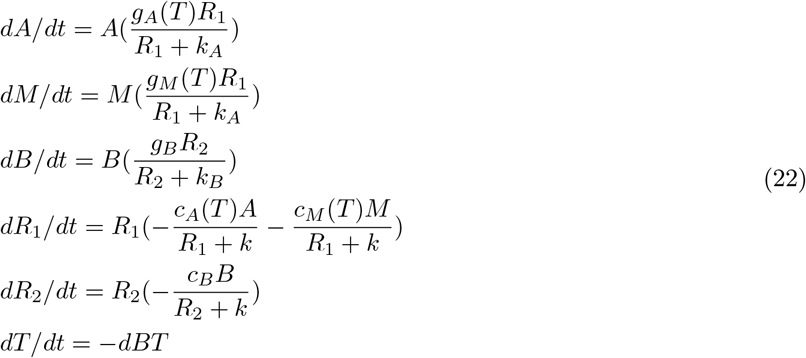

Taking into account such dynamics, we want to identify what mutants can establish and quantify the effect that detoxification has on their growth rates.

#### Detoxification-dependent selection across different mutants

Our approach consists of fixing an antibiotic concentration (*T* = 6.25 or 12.5, as in our experiment) and quantify how detoxification changes the selection acting on many randomly sampled mutations. The goal is test whether detoxification could change the distribution of fitness effects, explaining why we observed rescue in co-cultures but not in mono-cultures. Making use of the mechanistic model presented above, we generate thousands of mutants, simulate their growth under serial batch cycles and quantify their invasion selection. Each mutant is defined by its phenotype *g*_*M*_ = *g*_*A*_ + Δ_*g*_, *tol*_*M*_ = *tol*_*A*_ + Δ_*t*_ and *MIC*_*M*_ = *MIC*_*A*_ + Δ_*m*_, whereby the changes Δ are defined as follows:

- Δ_*g*_ = 0, 0.15, 0.3 or 0.45
- Δ_*t*_ ∼ *Normal*(*µ* = 0, *σ* = 10)
- Δ_*m*_ ∼ *Normal*(*µ* = 0, *σ* = 15)

When then simulate the dynamics of *A* in mono-culture or co-culture with *B*. During the first 3 transfers the antibiotic concentration is *T* = 0, then *T* = 6.25 or 12.5 in the remaining transfers. We assume that the mutant emerges at transfer 4 and study its dynamics afterwards. We measure selection acting on each mutant as the slope of its abundance over transfers, and not within transfers. This is because even if a mutant grows within a transfer it does not ensure survival over transfers due to dilution. Repeating this procedure for a total of 2000 randomly generated mutants, we show how detoxification by a second species *B* can increase the pool of mutants that would be able to invade and rescue the population from extinction. In other words, our simulations show that for the antibiotic concentrations under study, the presence of a detoxifying species can increase the fraction of beneficial mutations, thus affect a key parameter for evolutionary rescue: *U*_*s*_. This confirms our hypothesis that upon detoxification we need to take into account an additional parameter to explain the probability of rescue. We finally show an example of a mutant that can invade and rescue the population of *A* in co-culture, but fails to establish in mono-culture (Fig. **5**). The parameters used to produce the simulations for Fig. **5**a are as follows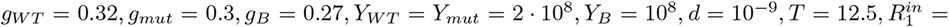,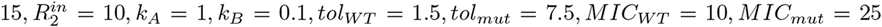where the wild type *WT* and the mutant *mut* represent two sub populations of species *A*.

### Mechanistic model of growth by resource consumption and detoxification by explicit betalactamase production

As an alternative to the previous model, here detoxification is performed by explicit enzymes, which is produced by the detoxifying species *B* and can accumulate over time. The dynamics follow:

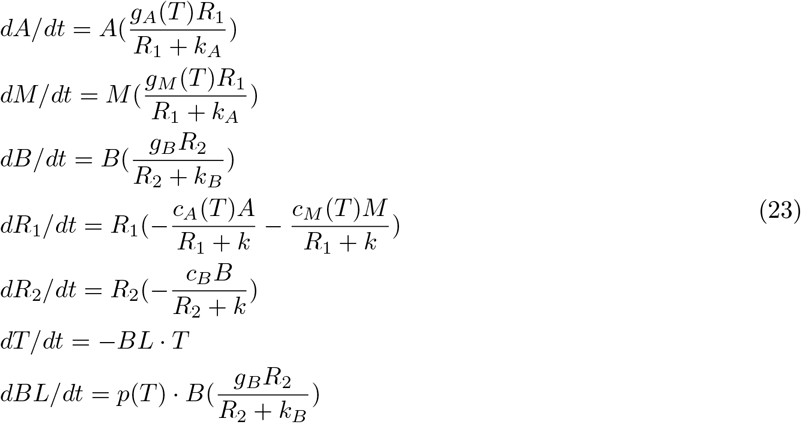

These dynamics are equivalent to the previous ones (eq.22), except the toxin concentration decays proportional to the beta-lactamase enzyme concentration (here *BL*), which in turn is produced upon growth via an antibiotic-dependent function *p*(*T*). This allows us to describe a scenario where the second species *B* does not produce *BL*, unless it encounters antibiotics. More specifically, we assume that:

- *p*(*T*) = 0 if *T* = 0, and *t > delay*
- *p*(*T*) = *P >* 0 otherwise

Assuming an activation delay of few hours, upon first encounter with antibiotics, this model can recapitulate the dynamics observed at high citrate high ampicillin (Fig. **4**b), whereby the susceptible population first drops and then slowly comes back, without any evolutionary process (Fig. S7). To obtain these dynamics we assume that the activation delay decreases over transfers (*delay* = 14*h*, 9*h*, 4*h* in the first 3 transfers with antibiotic and 0 afterwards) and *P* = 10^*−*9^ (the remaining parameters are as for the first mechanistic model). This mimics the scenario where beta-lactamase activation takes a few generations to synchronize throughout the entire population of *Ct*. While this parameter choice is completely arbitrary, the observation of delayed ecological rescue is conserved across different parameter regimes.

---

**Fig. S1:**
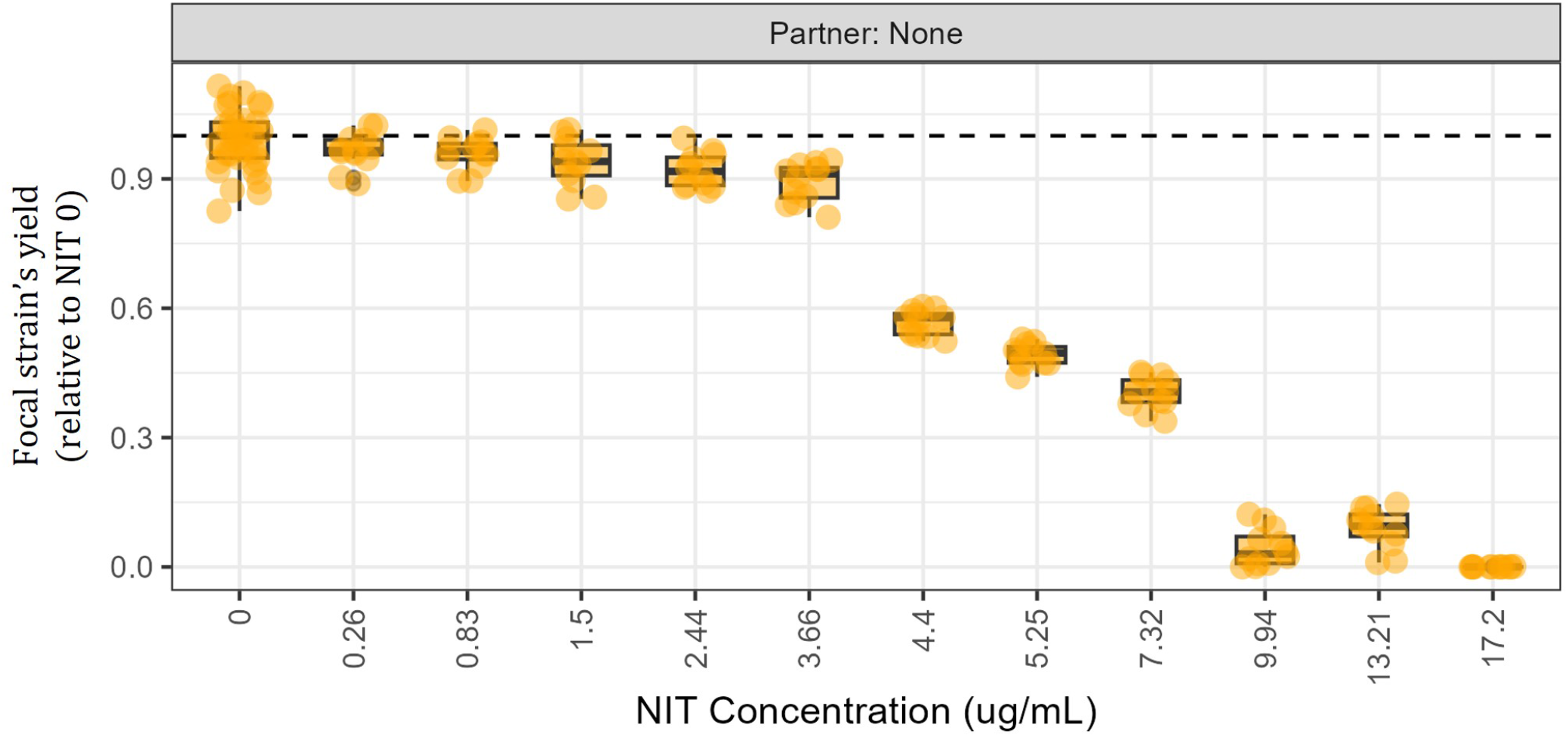
Sensitivity to nitrofurantoin. Max yield of the focal *E. coli* strain across nitrofurantoin (NIT) concentrations, relative to the NIT 0 case.

**Fig. S2:**
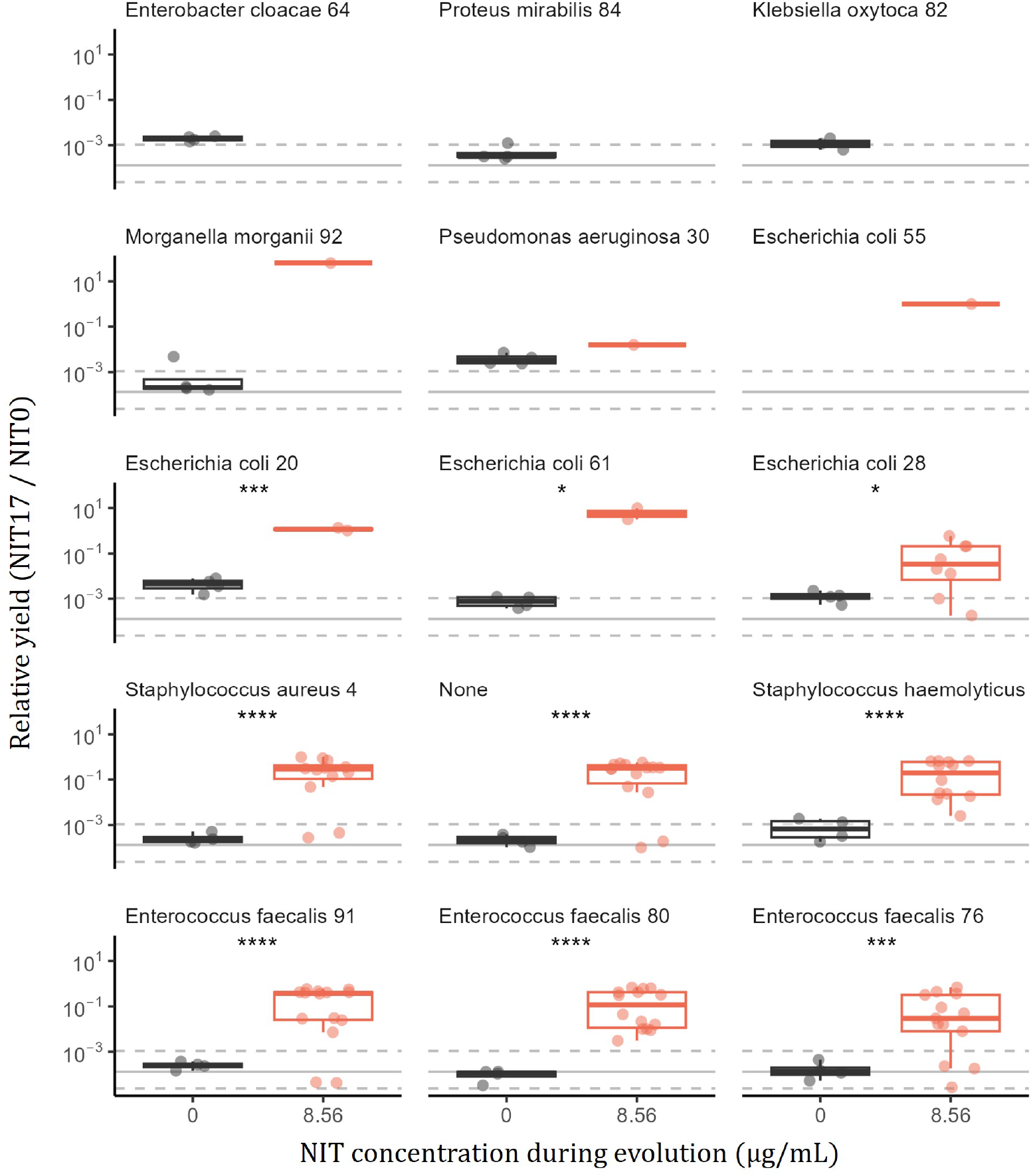
Evolved resistance across UTI partners. The data show the maximum observed luminescence under nitrofurantoin exposure (NIT 17 µg/mL) relative to the absence of it (NIT 0 µg/mL) of the focal *E. coli* populations evolved with (orange) or without nitrofurantoin (black) and in the presence of different UTI isolates. The solid gray line shows the mean relative yield of the ancestral *E. coli*, while the dashed gray lines indicate its minimum and maximum. Asterisks show significance level after t-test. The different number of replicates are due to the availability of survived populations.

**Fig. S3:**
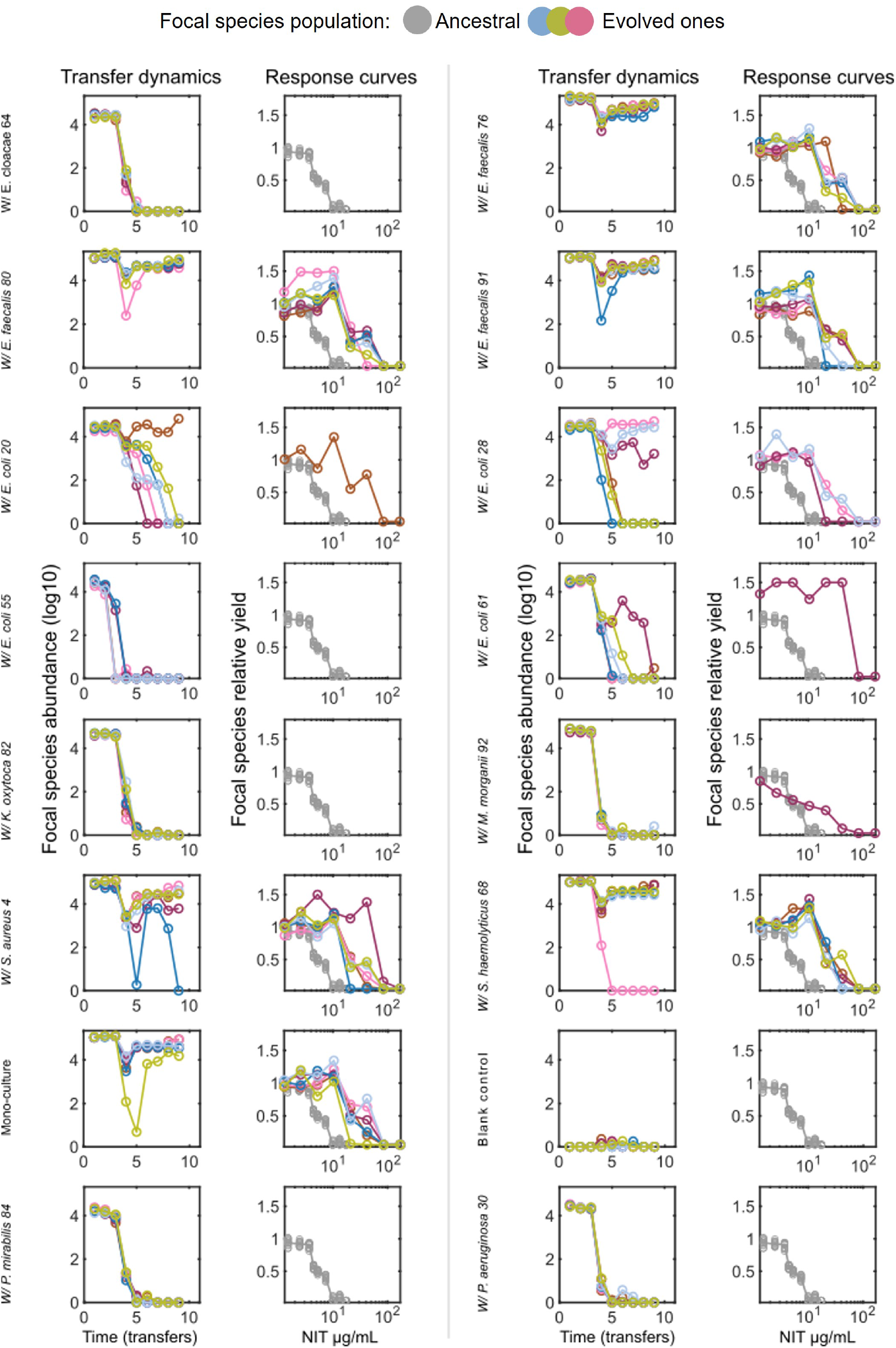
Evolutionary dynamics and dose-response curves of *E. coli* evolved populations. For each culture condition, six replicate cultures from a single experimental plate are shown. Columns 1 and 3 show the focal species abundance (in *log*_10_ of RLU, relative light units) across transfers. Columns 2 and 4 show the dose-response curves (DRCs), with drug concentration in µg/mL of nitrofurantoin (NIT), relative to no-drug conditions. Response values were capped between 0.05 and 1.5. DRCs were generated by outgrowth directly from the experimental plate, and population sizes are not standardized. Names on the side indicate the UTI isolate used in co-culture. Mono-culture: focal strain grown alone; Blank control: no strains added.

**Fig. S4:**
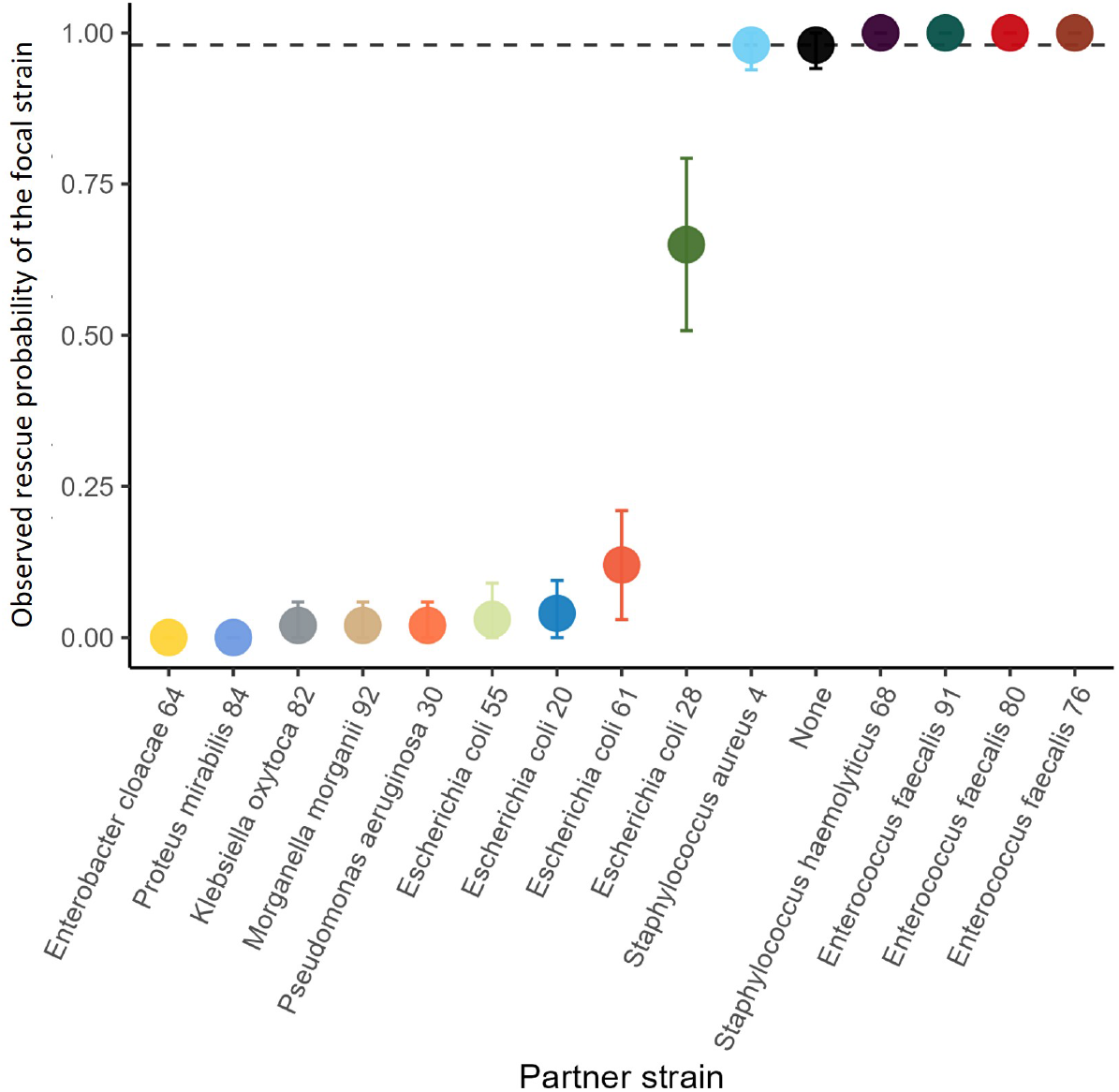
Observed rescue probability of the focal *E. coli* strain under the influence of different UTI partners. The rescue probability (*P*) is computed as the fraction of replicates that were found above the detection limit at the last transfer, over the total number of replicates (*N*). Data corresponds to the dynamics showed in Fig. **2**. The bars indicate the 95% confidence interval calculated as 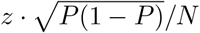, with *z* = 1.96.

**Fig. S5:**
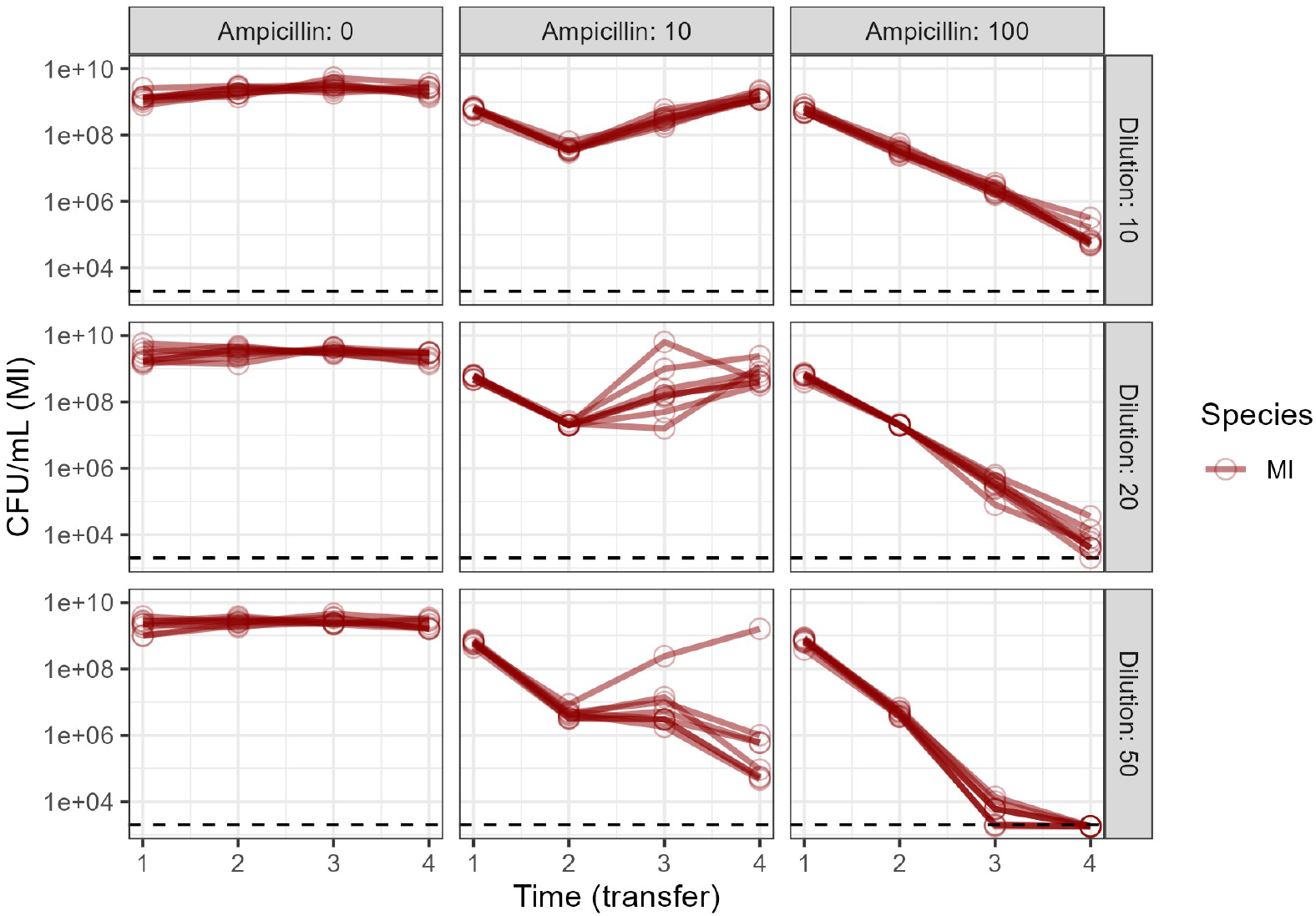
Effects of dilution rate and ampicillin concentration on the dynamics of *Ml*. Serial transfers of *Ml* occurring at different ampicillin concentrations (0, 10 or 100 µg/mL) and with different dilution factors (1:10, 1:20 or 1:50), 8 replicates per condition. Ideally we look for a regime where rescue is possible, but not too common. For this reason we picked the strongest dilution factor (1:50) and intermediate ampicillin (in the experiments we use 6.25 or 12.5 µg/mL).

**Fig. S6:**
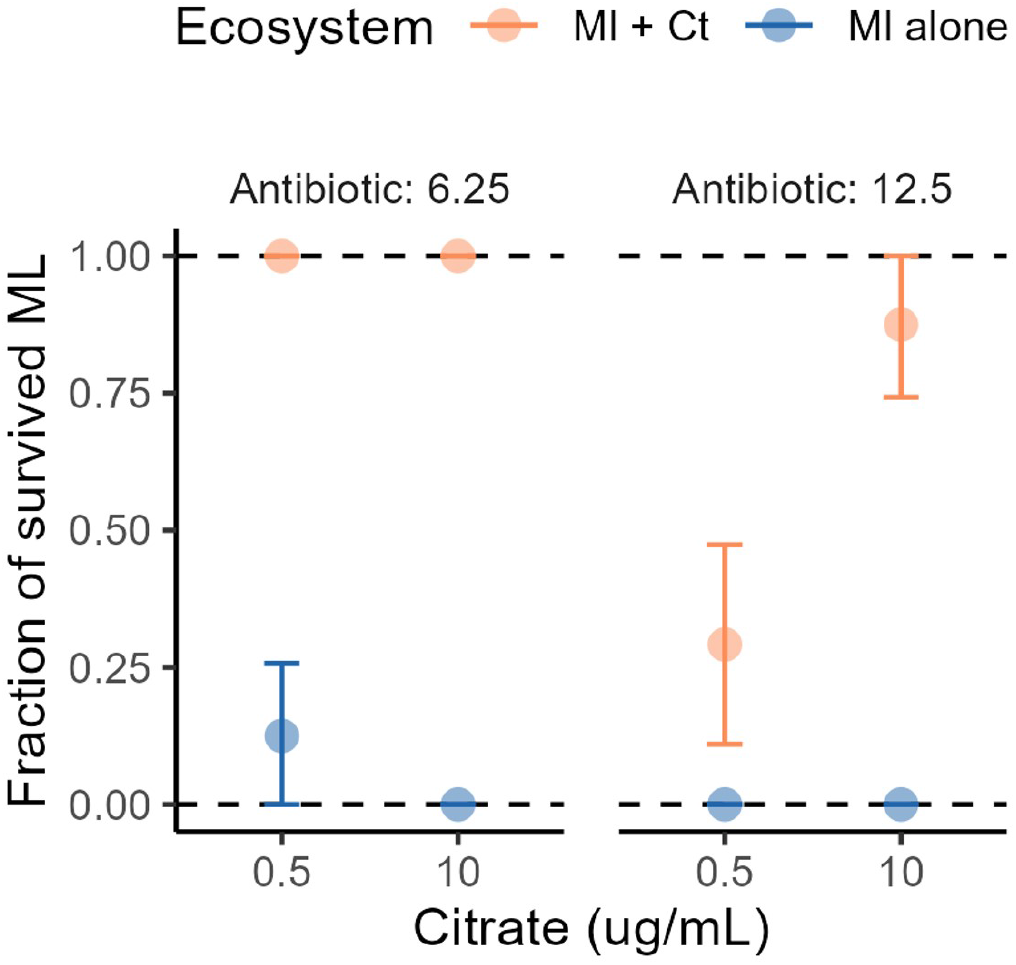
*Ml* survival probability across conditions. The survival probability (*P*) is computed as the fraction of replicates that were found above the detection limit at the last transfer, over the total number of replicates (*N*). Data corresponds to the dynamics showed in Fig. **4**b. The bars indicate the 95% confidence interval calculated as 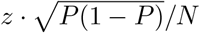, with *z* = 1.96.

**Fig. S7:**
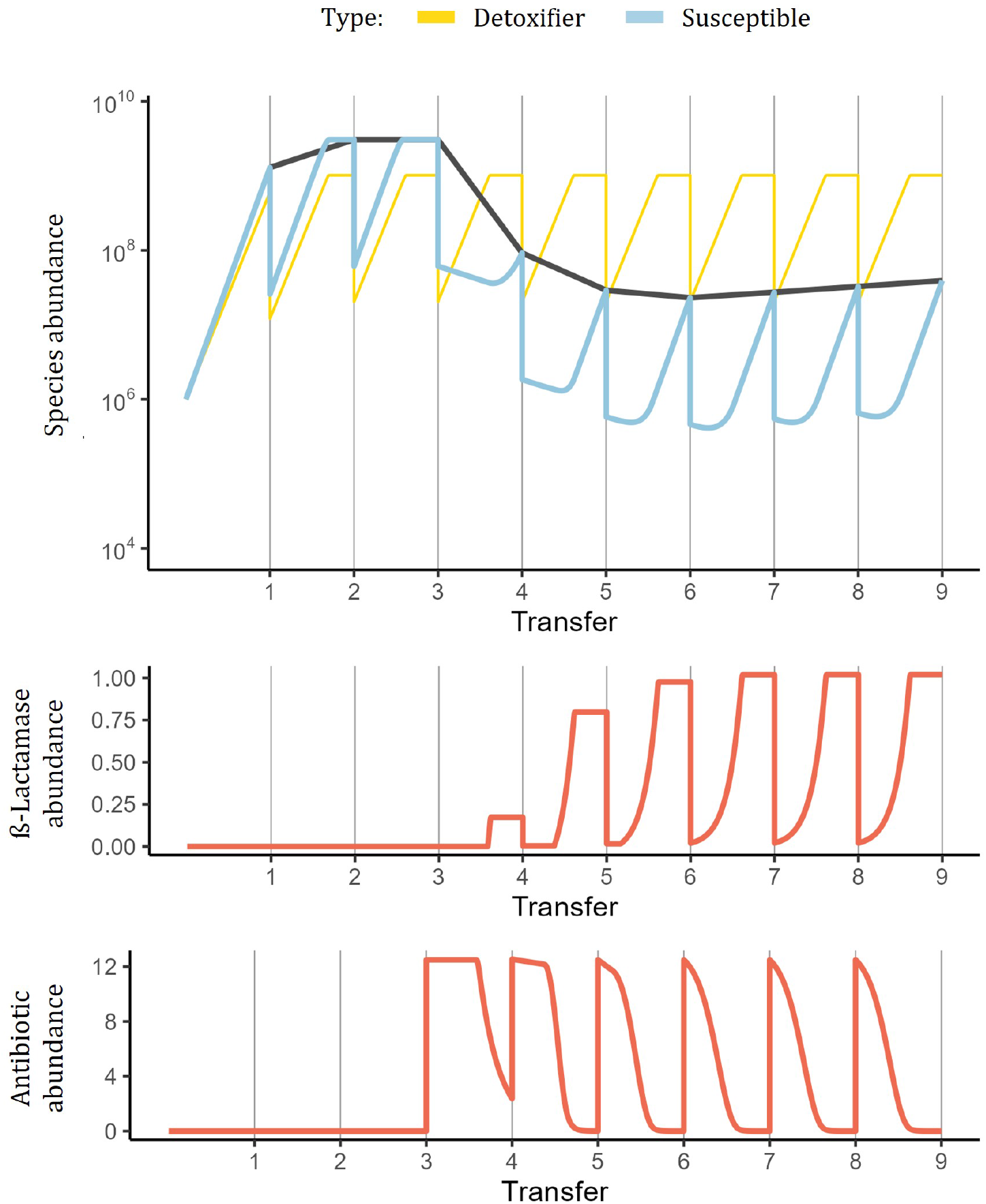
Delayed beta-lactamase production can recapitulate the delayed ecological rescue. Example simulation where a detoxifying and a susceptible species grow together over transfer, and experience antibiotic exposure after transfer 3. Here we assume a delay in the activation of beta-lactamase response, which causes a gradual accumulation of the beta-lactamase enzyme (middle panel), a delayed clearance of the antibiotic (bottom panel), and finally a delayed survival of the susceptible species. This example is coherent with our hypothesis of ecological rescue explaining the dynamics of *Ml* observed under strong-ampicillin-strong-detoxification regime (Fig. **4**)

**Fig. S8:**
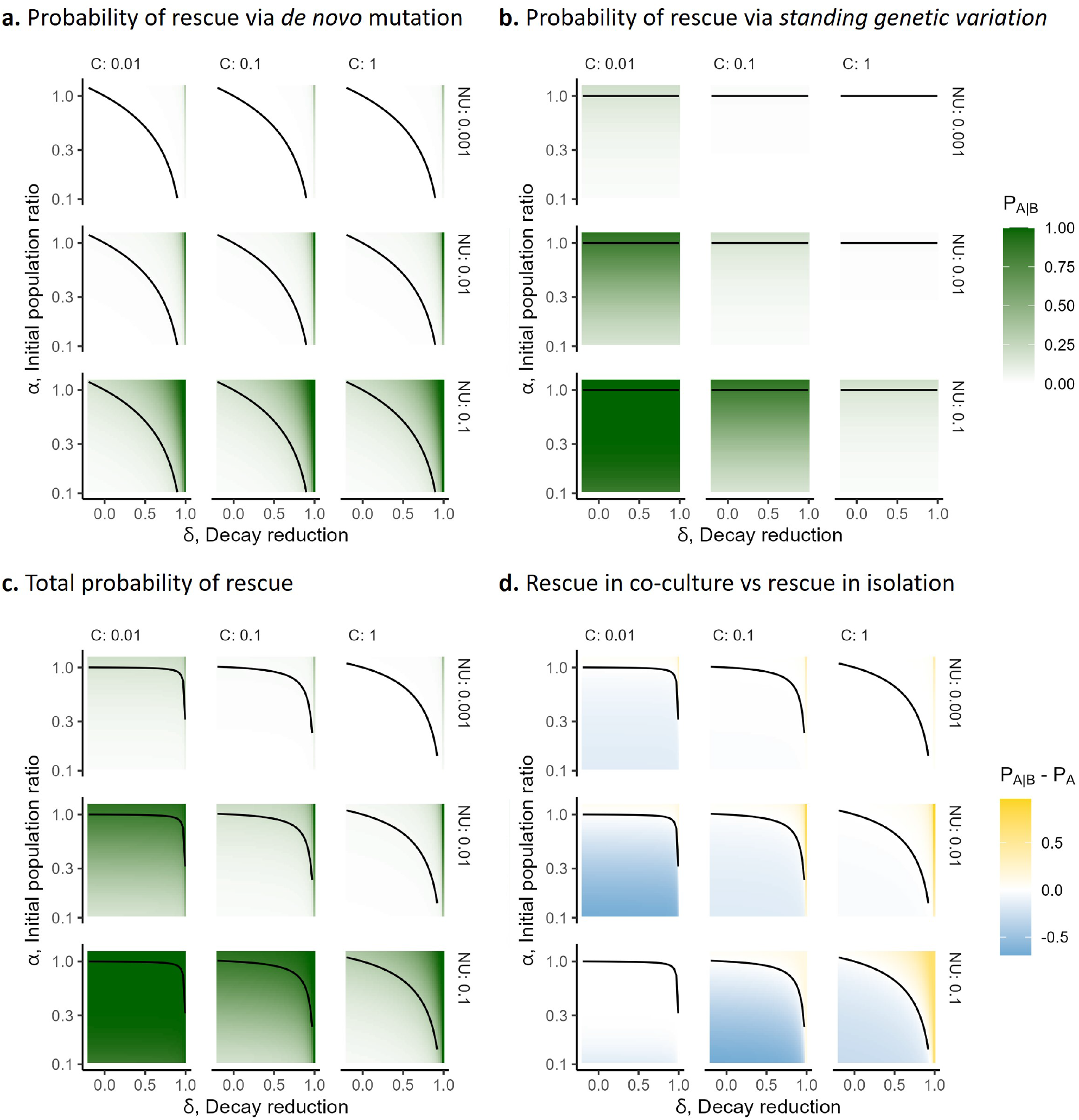
Evolutionary rescue probabilities under different conditions. **a,b,c)** Quantification of the absolute rescue probabilities via de novo mutations, standing genetic variation or both combined, as a function of the partner species effects on population size (*α*) or decay rate (*δ*). Across the column we vary the relative contribution of mutation rate to standing variation (*c* = *U*_*s*_*/p*_0_), while across rows we vary the effective input of mutations per generation (*N*, population size, *U* mutation rate). As expected the larger mutational inputs and larger variation (smaller *C*) increase rescue from standing variation, while only mutational input increases rescue via de novo mutations. **d** All together we show how even though the competition-balance is conserved across parameter conditions, the actual difference between rescue in co-culture and mono-cultures varies substantially.

**Fig. S9:**
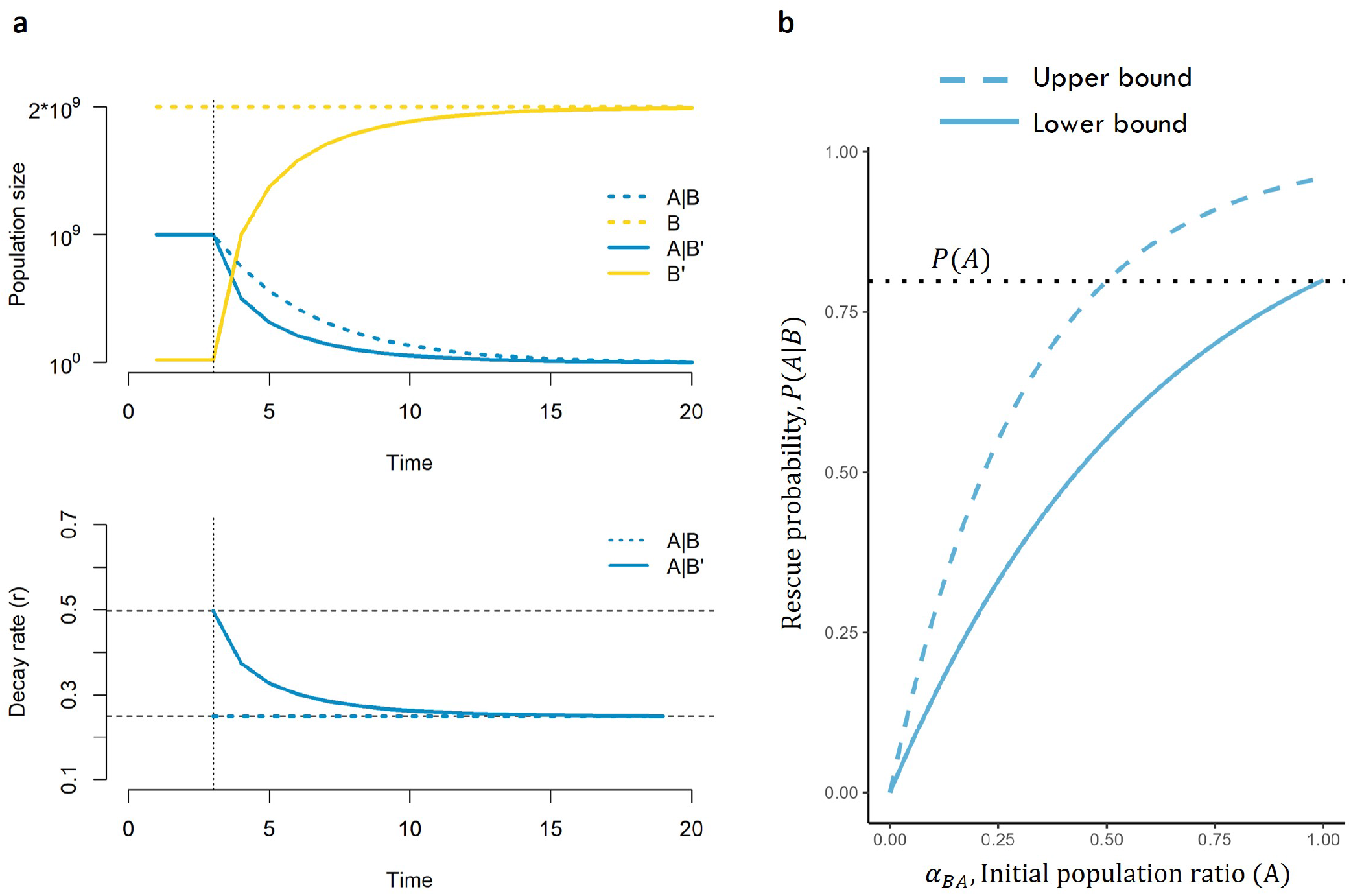
Rescue probabilities with density-dependent detoxification. **a)** Example of density-dependent dynamics where the abundance of the partner species *B* depends on the species *A* abundance (solid lines), compared to density-independent dynamics (dashed lines). Upper panel shows simulated dynamics where as soon as the toxins kicks in (vertical dotted line) species *A* declines, allowing species *B* abundance to increase. This leads to density-dependent decay rate of *A*, which starts faster and then decreases to finally approximate the density-independent case (bottom panel). **b** As a consequence the rescue probability of *A* decreases with the strength of competition between *A* and *B* (smaller *α*_*BA*_). When *α*_*BA*_ = 1 we retrieve the density-independent case studied in the main text.

**Fig. S10:**
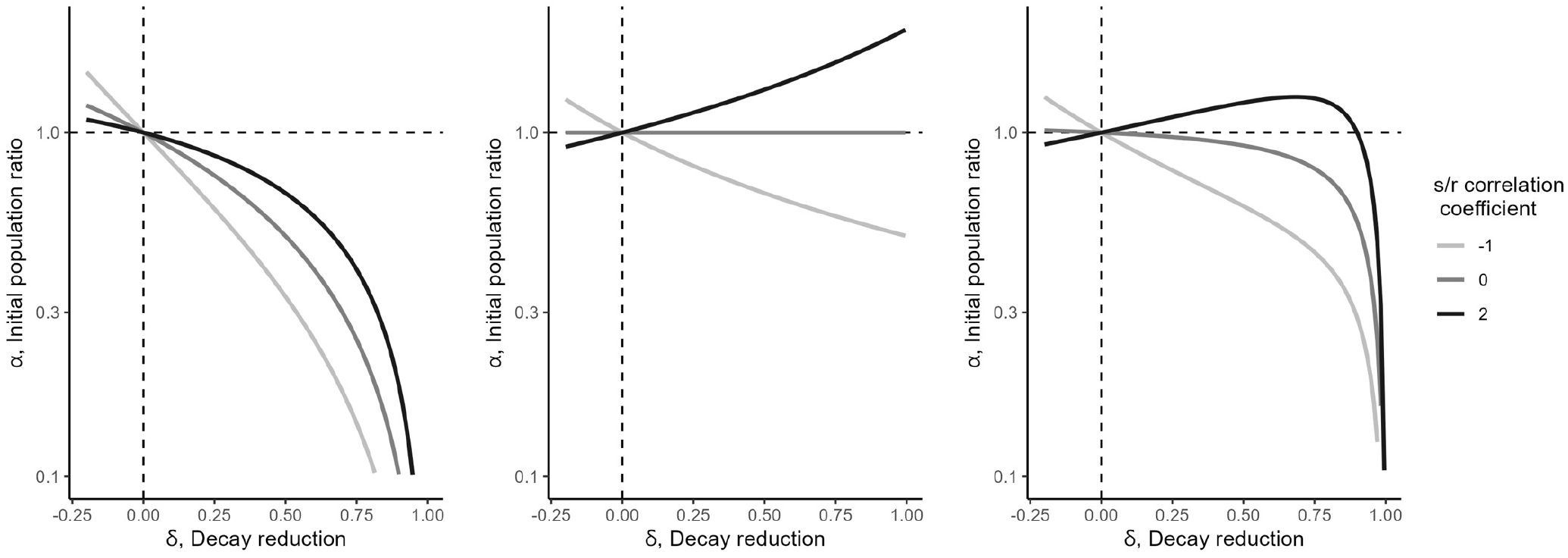
Competition-protection balance assuming a decay-selection correlation. Different shapes of the competition-protection balance based on the definition of decay and growth rates. As in Fig. **1**, above the curve we expect co-culture to favor rescue and vice versa below the curve. When the 2 are independent (correlation 0) we retrieve the case studied in the main text. Negative correlation means that the mutants grow faster when the susceptible type decays more slowly. This increases the parameter region where having a second species favors rescue. Contrarily, positive correlation means that the mutants grow slower when the susceptible type also decays more slowly. This reduces the parameter region where having a second species favors rescue.

## Notes

### Competing Interest Statement

The authors have declared no competing interest.

